# Metabolic reprogramming promotes *Staphylococcus aureus* serum resistance during bacteraemia

**DOI:** 10.64898/2026.01.14.699454

**Authors:** Samuel Fenn, Ruth C. Massey

## Abstract

*Staphylococcus aureus* is a leading cause of bloodstream infections causing an estimated 300,000 deaths worldwide. Using a functional genomics approach, our group previously identified that adaptation of *S. aureus* to human serum is polygenic, with a clinically occurring non-synonymous mutation (V76I) in dihydrolipoamide dehydrogenase (*lpdA1*/*pdhD*) improving bacterial survival. In this work we establish that improved serum survival of strains expressing PdhD V76I is underpinned by enhanced resistance to host-derived antimicrobials including antimicrobial peptides and host-defence fatty acids which are prevalent in the bloodstream. Here we demonstrate that the PdhD V76I variant has enhanced diaphorase activity, with both clinical and laboratory strains expressing this variant recycling NADH to NAD^+^ without using respiration, leading to a reduction in membrane potential. This conferred persister / small colony variant (SCV) like phenotypes on strains expressing PdhD V76I including increased resistance to gentamicin, hydrogen peroxide, LL37, HNP-1 and arachidonic acid. However, strains which utilise PdhD V76I do not display growth defects typical of persisters and SCVs, with reduced NADH accumulation in serum leading to enhanced glycolysis/TCA cycle activity, enhancing bacterial replication in human serum. Whilst establishment of SCV and persister populations is a key survival strategy in the bloodstream, this work demonstrates how intermediate phenotypes can also be effective at promoting survival in this hostile environment. This highlights the heterogenous nature of *S. aureus* adaptation to the host-environment, with an improved fundamental understanding of these processes required to allow for the development of novel therapeutics which target this process of host-adaptation.

**IMPORTANCE:** The establishment of a bloodstream infection requires the bacteria to be able to withstand the antibacterial factors found there. As one of the major global causes of these types of infections, *S. aureus,* utilises many diverse strategies to survive and replicate in the bloodstream, which can include modifications to either the cell envelope, or to core metabolic processes such that the host’s immune features are withstood. While extreme adaptations to metabolism can lead to the major growth defects associated with persisters and small colony variants, here we describe a clinically relevant adaption where metabolism was altered such that the bacteria could survive and replicate in serum, but where the growth characteristics of the bacteria were not affected. Mutation of the gene encoding the dihydrolipoamide dehydrogenase enzyme (*lpdA1*/*pdhD*) enhanced it diaphorase activity such that it could recycle NADH to NAD^+^ without using respiration, which led a reduction in membrane potential that conferred resistance to gentamicin, hydrogen peroxide, LL37, HNP-1 and arachidonic acid and human serum, but did not affect growth. This work uncovers a novel mean of adaptation to the bloodstream by *S. aureus* and highlights how it plasticity underlies its success as a major human pathogen.

## INTRODUCTION

*Staphylococcus aureus* is an important human pathogen responsible for over 1 million deaths a year (1). This opportunistic pathogen associates with humans as a commensal, with transition to a pathogenic state responsible for a wide range of diseases ranging from soft tissue to urinary tract infections (2). The most dangerous of these diseases is bacteraemia, with *S. aureus* bloodstream infections killing in 20-30% of cases causing an estimated 300,000 deaths per year (3–5). *S. aureus* survival and persistence in the bloodstream is a key component to progression of this disease. This opportunistic pathogen gains access to the bloodstream through initial colonisation of other sites or through intravenous medical devices. Once in the bloodstream, this pathogen can disseminate to all body parts with the resulting sepsis often leading to multiple-organ failure (2,6). The ability of this pathogen to form biofilms, establish intracellular reservoirs, small-colony variants, and persister cells increases antimicrobial tolerance, complicating treatment of *S. aureus* once an infection has embedded (7–9).

Initial establishment of an infection at a host site is mediated by an extensive arsenal of virulence factors and defensive mechanisms. Toxins, proteases and lipases secreted by *S. aureus* promote local cellular damage and liberate nutrients from the host environment (10). Whilst immune evasion strategies such as production of Spa/Sbi block antibody binding, with proteins such as Eap, SCIN and Ecb inhibiting complement activation (6). This enables *S. aureus* to evade the action of the host immune system and persist in the host environment. Infection of the bloodstream by *S. aureus* is heavily dependent on defensive mechanisms, with invasive diseases such as bacteraemia or lung infections generally accepted to result in downregulation of virulence factor production (11,12). These defences are required as the bloodstream is a heavily protected niche with multiple components exhibiting anti-staphylococcal action, such as antimicrobial peptides (AMPs) and host-defence fatty acids (HDFAs) (13–16). The ability of *S. aureus* to overcome these host defences is critical to establishment of a bloodstream infection, with this organism capable of adapting and evolving in-host to resist these anti-staphylococcal factors.

One key mechanism of adaptation in hostile environments is modulation of metabolism (17). Invasion of the bloodstream naturally leads to reduced growth capacity of *S. aureus*, with the innate immune system locking away key metabolites such as iron and copper by chelating them from the environment in a process known as nutritional immunity (18,19). This leads to respiratory dysfunction and results in naturally higher levels of tolerance by host-derived antimicrobials. *S. aureus* is also capable of producing small-colony variants (SCVs), with mutations which negatively impact the respiratory chain (*hemB*, *menD*) commonly identified *in vivo* (7,20). These SCVs exhibit slow growth due to reduced metabolic activity, increased antimicrobial tolerance and reduced toxicity, leading to reduced antibacterial activity of both the immune system and antibiotics (7). Similarly, alterations in central carbon metabolism also reduce susceptibility to host and exogenous antimicrobials, with interruption of the TCA cycle promoting survival in these hostile host environments (21,22). Once these stresses are removed, SCVs can revert to wild-type or acquire suppressor mutations which enable maintenance of resistance whilst improving replication of *S. aureus*.

This work builds upon previous work conducted in our group detailing the application of a genome wide association study (GWAS) to identify polymorphisms associated with altered survival in human serum (23). One of the novel effectors identified in this study was the dihydrolipoamide dehydrogenase *lpdA*/*pdhD*. This enzyme re-oxidises dihydrolipoamide to lipoamide on the E2 subunit of the pyruvate dehydrogenase complex. This enables the pyruvate dehydrogenase complex to continue producing acetyl CoA from pyruvate, feeding into both the TCA cycle and fatty acid synthesis (24). A Valine76Isoleucine amino acid substitution in PdhD was identified to drastically improve survival of *S. aureus* in serum, whilst *pdhD* deletion resulted in reduced resistance to serum suggesting a gain of function mutation had occurred (23). In this work we demonstrate that the V76I mutation metabolically reprogrammes *S. aureus* through improved redox cycling of NADH to NAD^+^. This confers some SCV like phenotypes without compromising the growth of *S. aureus*, with strains expressing the PdhD V76I mutant demonstrating increased resistance to host and exogenous antimicrobials. As a result, these strains can persist and grow in human serum, demonstrating that establishment of SCVs or persister cells is not the only pathway to survival of *S. aureus* in the bloodstream.

## RESULTS

### Dihydrolipoamide dehydrogenase mutation increases *S. aureus* serum resistance

Human serum contains a diverse array antimicrobial compounds which work synergistically to maintain blood sterility (13–16). In a previously performed genome-wide association study screening for genes which impact survival in serum a V76I amino acid substitution in PdhD was associated with improved survival in serum however the physiological basis was not explored for the improved survival (Figure 1A). To isolate the effect of the V76I substitution, we created a pCN34 based complementation plasmid with *pdhD* transcription under control of the native *pdhA* promoter (25). To construct both wild-type and V76I allele expressing complementation plasmids a sequential approach was taken. The *pdhA* promoter region was amplified from the CC30 wild-type strain MRSA252 and cloned into pCN34. Next the *pdhD* sequences from MRSA252 and ASARM161 (PdhD V76I) were amplified and inserted downstream of the P*_pdhA_* promoter creating plasmids p*pdhD* (wild type *pdhD*) and pSNP (mutant *pdhD*). These vectors were then electroporated into the MRSA strain JE2, and two MSSA strains SH1000 and Newman harbouring the *pdhD*::tn mutation, and screened for their ability to survive in serum. As previously reported, strains lacking *pdhD* demonstrate reduced survival in human serum, with both JE2, SH1000 and Newman *pdhD*::tn mutants exhibiting a 43, 52% and 50% reduction in colony forming unit (CFU) recovery respectively (Figure 1B). This demonstrates that the *pdhD*::tn phenotype is not JE2 or MRSA specific. Restoration of the wild type *pdhD* allele restores serum resistance to wild-type levels, whilst complementation with PdhD V76I further increases *S. aureus* CFU recovery and fosters active growth in serum after 2 hours of incubation for both JE2 and SH1000 isogenic strains (Figure 1C and S1). This confirms the effect of the V76I mutation identified during the serum resistance GWAS (23).

**Figure 1.**
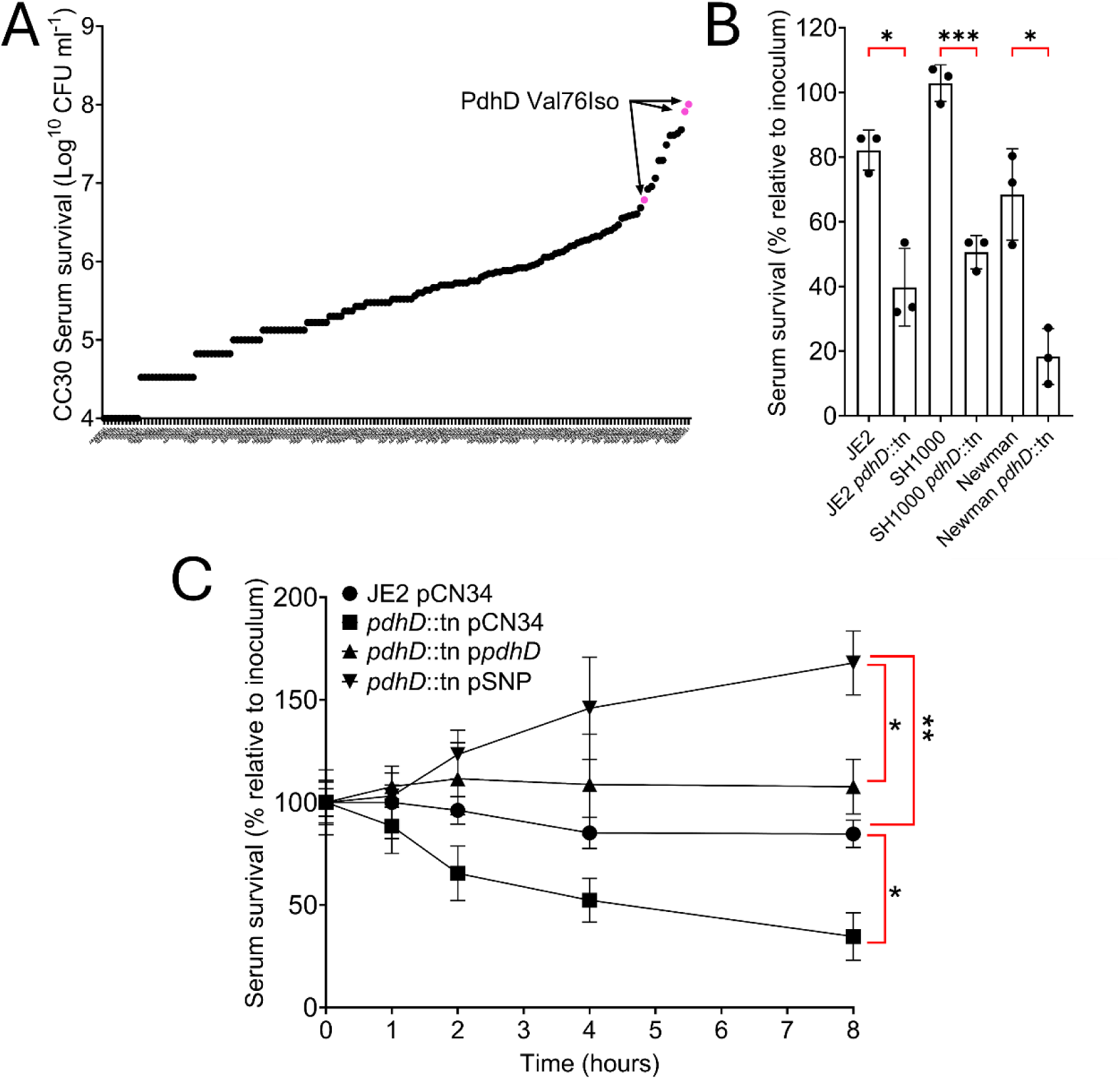
Expression of the PdhD V76I variant encourages enhances survival and encourages growth of *S. aureus* in human serum. (A) Visualisation of PdhD V76I amino acid substitution identified during a genome wide association study as improving *S. aureus* serum survival. (B) Loss of *pdhD* compromises *S. aureus* serum survival in both MRSA (JE2) and MSSA (SH1000/Newman) genetic backgrounds. (C) Survival of S. aureus *pdhD* mutant and complemented strains in 25% human serum. Points represent average of three experiments in A and C whilst bars represent the mean value in B. Error bars represent standard deviation. Significance determined as *<0.05, **<0.01, ***<0.001, calculated by one-way ANOVA with Dunnett’s T3 (B) and two-way ANOVA with Tukey’s (C) multiple comparisons test.

### Strains expressing PdhD V76I are more resistant to host derived antimicrobials

Our group previously demonstrated that loss of *pdhD* compromised survival in serum (23), whilst we have now established that the V76I substitution in PdhD increases serum resistance (Figure 1C). To establish if this mutation provides a general protective effect or is specific to a subset of serum-derived antimicrobials we screened susceptibility to cationic antimicrobial peptides and host defence fatty acids, with these antimicrobials acting a barrier to establishment of bloodstream infections.

Strains lacking *pdhD* demonstrate enhanced susceptibility to the antimicrobial peptide LL37 and HNP-1, with a 1.4 and 1.7 log reduction in CFU/mL recovery as compared to wild type respectively (Figure 2A-B). This is in line with the enhanced killing of *pdhD* mutants in serum (Figure 1C). Complementation with the native *pdhD* allele restores *S. aureus* resistance to LL37 and HNP-1, whilst complementation with the V76I PdhD variant provides a significant increase in protection against both antimicrobial peptides (Figure 2A-B). A similar trend is observed when testing killing of these strains by arachidonic acid (AA) with loss of *pdhD* sensitising *S. aureus* to AA, whilst complementation with the V76I substitution enhances *S. aureus* resistance when compared to native *pdhD* complementation (Figure 2C). Clinical strains from the original serum survival screen expressing PdhD V76I were then screened for susceptibility to antimicrobial peptides and host defence fatty acids. *S. aureus* isolates harbouring this mutation demonstrate increased resistance to LL37, HNP-1 and AA when compared to a random selection of CC30 isolates (Figure S2). This confirms the biological significance of the naturally occurring V76I mutation and demonstrates that this allele provides a generic protective effect against serum derived antimicrobials. Increased resistance to these antimicrobials enables strains expressing the PdhD V76I to quickly initiate replication in serum, explaining identification of this allele in our previously performed GWAS (23).

**Figure 2.**
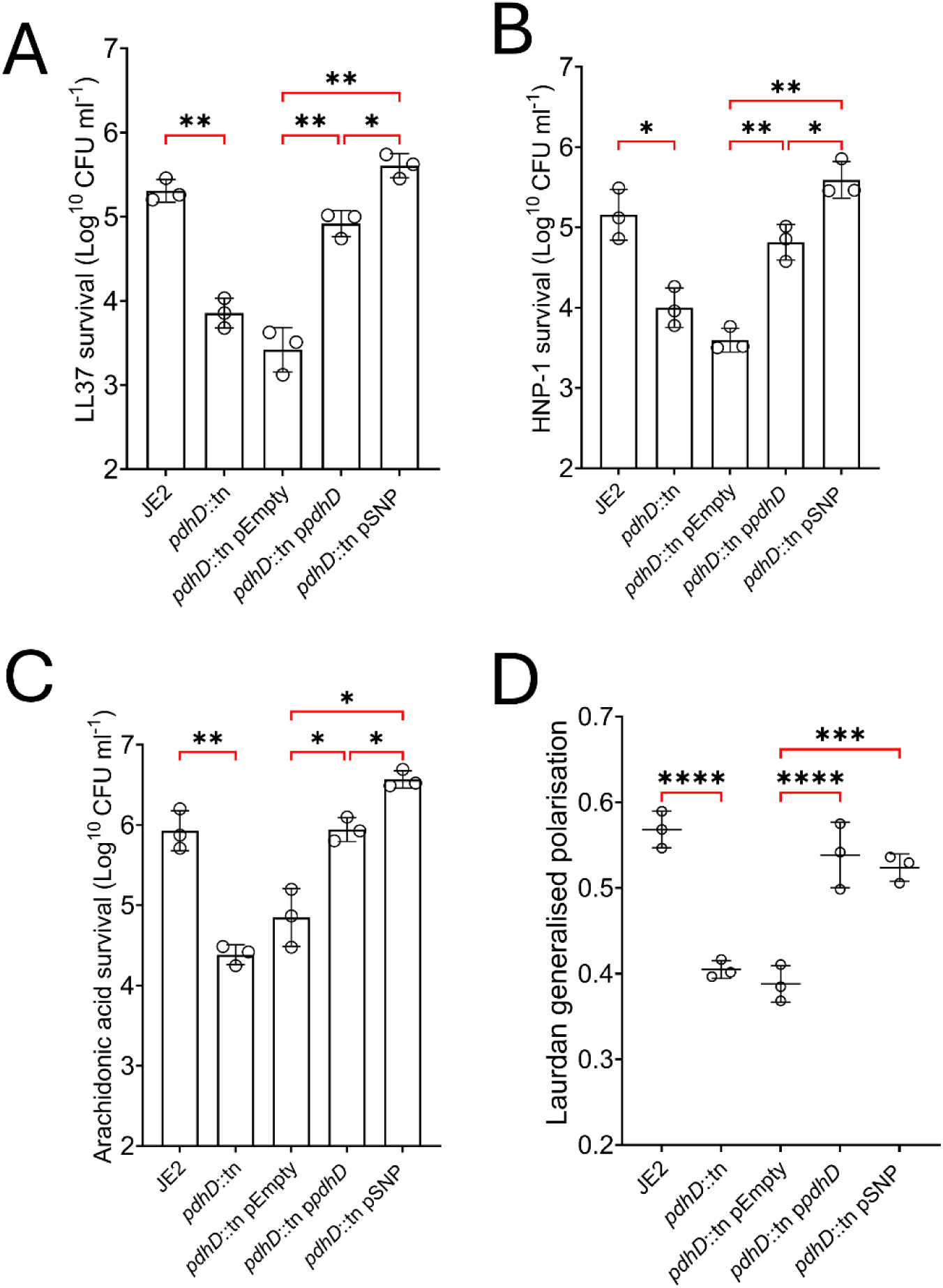
– PdhD V76I increases CAMPs and HDFA resistance without impacting cytoplasmic membrane fluidity. *S. aureus* strain killing assays with LL37 (A), HNP-1 (B), and Arachidonic acid (C). (D) Membrane fluidity assessed with Laurdan dye. LL37 and HNP-1 killing assays conducted in PBS at 5 µg mL^-1^ for 4 hours with a starting inoculum of 1x10^6^ cells. Arachidonic acid killing assays conducted in TSB at 200 µM and 2.5 µg mL^-1^ for 2 hours with a starting inoculum of 1x10^7^ cells. Laurdan dye added at 10 µM for 10 minutes followed by washing with emissions detected at 440nm and 490nm used to infer the fluidity of the plasma membrane. Each dot represents and individual experiment with a minimum of three technical replicates technical replicates. Bars and lines represent mean values, error bars the standard deviation. Significance determined as *<0.05, **<0.01, ***<0.001, ****<0.0001 calculated by one-way ANOVA with Sidak’s multiple comparisons test.

Acetyl-CoA produced by the pyruvate dehydrogenase complex feeds into both the TCA cycle and straight chain fatty acid synthesis, with the latter dictating phospholipid membrane composition and fluidity. Loss of *pdhA* and the subsequent reduction in the straight chain fatty acid synthesis, has been reported to increase membrane fluidity of *S. aureus* (24). This renders strains more susceptible to attack by membrane active antimicrobials such as LL37, HNP-1 and AA. Alterations in membrane fluidity could impact susceptibility to these antimicrobials altering survival in serum. To determine if the membranes of strains expressing PdhD V76I are altered, we employed Laurdan labelling as a direct measure of membrane fluidity (26). Complementation with wild type *pdhD* and *pdhD* V76I restored membrane fluidity to the same extent (Figure 2D), confirming that alterations in membrane fluidity are not responsible for the increased resistance to LL37, HNP-1 and AA of the PdhD V76I expressing strains.

### The *pdhD* SNP allele increases *S. aureus* hydrogen peroxide and gentamicin resistance without inducing a small colony variant phenotype

Acetyl-CoA produced by the pyruvate dehydrogenase complex is the main source of fuel for the TCA cycle, reacting with oxaloacetate to form citrate. Mutants of the pyruvate dehydrogenase complex adopt a small colony variant (SCV) phenotype with loss of *pdhA* and *pdhB* shown to compromise replication of *S. aureus* (24). Visual examination of individual colonies of the isogenic wild type and mutant pairs, and the clinical strains containing the V76I mutation verified that they were all normal in size, however, to ensure this mutation does not alter growth kinetics we performed growth curves in both TSB and RPMI-1% casamino acids (RPMI-C). As expected, strains deficient for *pdhD* demonstrate reduced growth in both TSB and RPMI-C (Figure 3A-B). Complementation with the native *pdhD* allele and *pdhD* V76I restored growth kinetics to wild-type levels in both TSB and RPMI-C suggesting there is no growth defect associated with the V76I mutation (Figure 3A-B). SH1000 isogenic strains demonstrate the same pattern whilst growth kinetics for the clinical strains encoding the V76I mutation in PdhD demonstrate no defect in TSB or RPMI-C (Figure S3).

**Figure 3.**
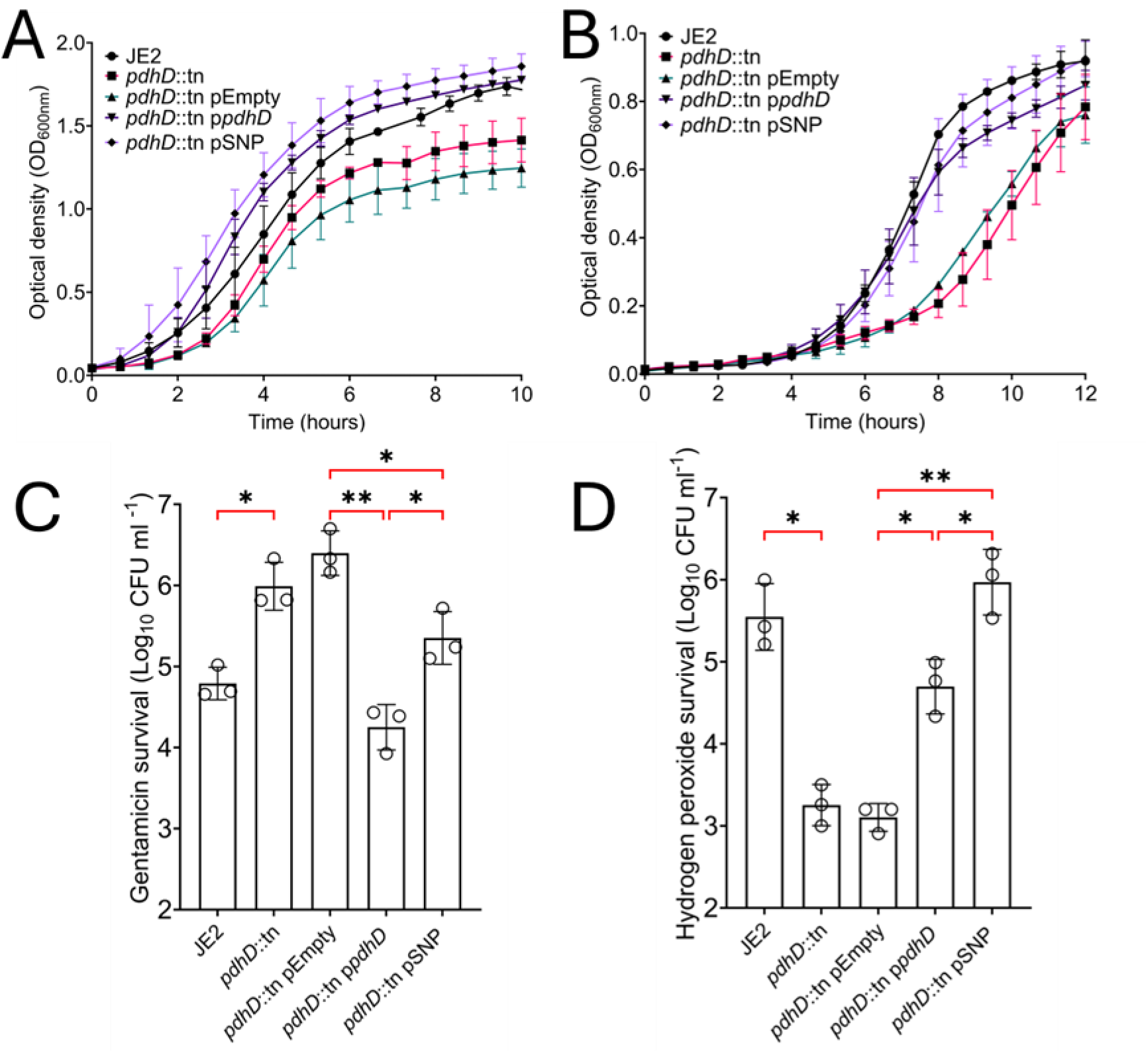
PdhD V76I mutation provides protection against gentamicin and hydrogen peroxide without inducing a small colony variant phenotype. (A-B) Growth curves of *pdhD* mutant and complements in TSB and RPMI-C respectively. S. aureus killing assays with gentamicin (C) and hydrogen peroxide (D). For growth kinetics points represent the average of three independent experiments with a minimum of four technical replicates per experiment. Gentamicin and hydrogen peroxide killing assays conducted with an initial inoculum of 1x10^7^ cells. Gentamicin killing conducted in TSB at 5 µg mL^-1^ for 2 H, hydrogen peroxide killing conducted in PBS at 20 mM for 90 minutes. Points represent individual experiments with a minimum of three technical replicates; bars represent the mean values with error bars standard deviation. Significance determined as *<0.05, **<0.01 calculated by one-way ANOVA with Sidak’s multiple comparisons test.

Increased hydrogen peroxide and gentamicin resistance are hallmarks of SCV in *S. aureus*. However, these phenotypes can also be used to screen for metabolic modifications, with killing by hydrogen peroxide and gentamicin dependant on a functional electron transport chain (27,28). Typical of SCVs, a *pdhD* mutant demonstrated increased resistance to gentamicin, whilst atypical of SCVs, loss of *pdhD* sensitises *S. aureus* to hydrogen peroxide mediated killing (Figure 3C-D). Complementation with native *pdhD* restores the gentamicin and hydrogen peroxide killing phenotypes back to wild type. However, restoration of the *pdhD* SNP allele only partially restores gentamicin killing whilst simultaneously increasing the ability of S. aureus to resist hydrogen peroxide when compared to complementation with the native *pdhD* allele (Figure 3C-D). These phenotypes were confirmed with MIC assays where *pdhD* V76I complementation increases resistance to gentamicin and hydrogen peroxide without impacting the MIC of other tested antimicrobials in both JE2 and SH1000 backgrounds (Table S1). Despite this elevated resistance PdhD V76I containing strains do not possess a growth defect under tested conditions ruling out the SCV phenotype. Given the role of *pdhD* in central carbon metabolism and associated increases in hydrogen peroxide/gentamicin resistance, it is likely that the increased serum survival exhibited by this SNP is metabolically mediated.

### PdhD V76I demonstrates increased diaphorase activity

Given the increased resistance to gentamicin and hydrogen peroxide we examined whether this mutation is associated with defective pyruvate dehydrogenase function. An interruption in central carbon metabolism would reduce TCA and electron transport chain activity, elevating tolerance to multiple antimicrobials. PdhD catalyses two reactions, a dihydrolipoamide dehydrogenase (DLD) reaction which regenerates the essential lipoate cofactor in the pyruvate dehydrogenase complex, and a diaphorase (DP) reaction which oxidises NADH to NAD^+^ using an external electron acceptor (Figure 4A) (29). To understand the biochemical effect of the V76I on PdhD catalytic function we purified both variants for enzymatic activity assessment.

**Figure 4.**
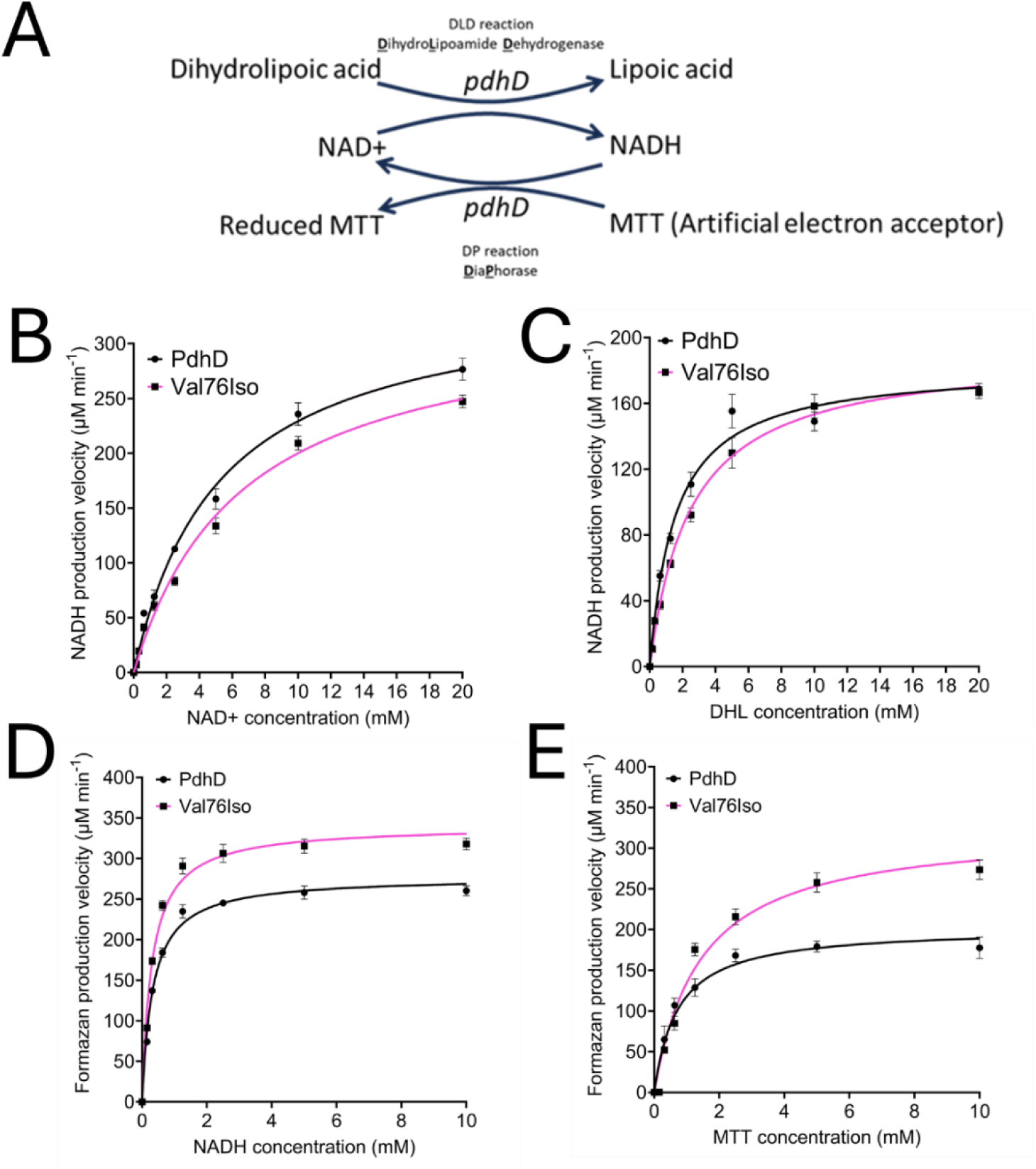
PdhD V76I mutation causes a minor reduction in dihydrolipoamide dehydrogenase activity whilst increasing diaphorase activity. (A) Schematic diagram outlining reaction schemes for dihydrolipoamide dehydrogenase (DLD) and Diaphorase (DP) assays. DLD assays performed with variable NAD^+^ (B) and dihydrolipoic acid (C). DP assays performed with variable NADH (D) and MTT (E). All reactions conducted in 0.1 M Potassium phosphate buffer pH 7.2 at 30^°^C. DLD assay measured through appearance of NADH at A_340nm_. DP assay measured through production of formazan product at A_570nm_. Points represent mean of three independent experiments with error bars indicating standard deviation. At least two technical replicates were conducted per experiment Reaction velocities for Michaelis-Menten kinetics calculated using GraphPad prism.

The enzymatic kinetic parameters of the DLD reaction were characterised using a reaction consisting of NAD^+^, dihydrolipoic acid (DHL) and PdhD with production of NADH measured spectrophotometrically (Figure 4A). Initial reaction velocities were determined at varying substrate concentrations for both NAD^+^ (Figure 4B) and DHL (Figure 4C). These values were used to determine kinetic parameters by non-linear regression and Michaelis-Menten kinetics. This revealed that the V76I mutation results in a small decrease in substrate affinity (K_m_) and catalytic efficiency (K_cat_/K_m_) for NAD^+^and DHL (Table 1A), indicating that this mutation reduces the ability of PdhD to bind DHL/ NAD+ and regenerate lipoate. We next sought to characterise diaphorase (DP) activity of this variant using a reaction consisting of NADH alongside the artificial electron acceptor MTT which is reduced to form a purple formazan product, enabling kinetic tracking of DP activity (Figure 4A). Initial reaction velocities were determined at varying concentrations of NADH (Figure 4D) and MTT (Figure 4E), with kinetic parameters determined as performed for the DLD assay. PdhD V76I demonstrated enhanced diaphorase activity when using NADH (Figure 4D and Table S2) and MTT (Figure 4E and Table S2) as substrates. The effect of this mutation on DP activity is more drastic than that observed for the DLD assay (Figure 4B-C) with the V76I form of PdhD achieving higher maximum reaction rates (V_max_); elevated number of substrate molecules a single enzyme can convert into product per second (k_cat_); and increased catalytic efficiency (k_cat_/K_m_) when using NADH and MTT as substrates (Table S2).

### Increased diaphorase activity reduces *S. aureus* intracellular NADH accumulation

Based on the increased diaphorase activity exhibited by PdhD V76I we hypothesized that this enzyme is contributing to NADH reoxidation to NAD^+^. Maintenance of the NADH: NAD^+^ ratio is essential to metabolic function of *S. aureus* with NAD^+^ serving as an essential cofactor in the TCA cycle, and NADH used by type-II NADH dehydrogenases to reduce menaquinone for use in the electron transport chain (ETC) (30). High NADH: NAD^+^ ratios are indicative of reductive stress and lead to inhibition of the TCA cycle and respiration, whilst the inverse is true for low NADH: NAD^+^ ratios. The ability of *S. aureus* to alter its metabolism in response to stress is pivotal to its ability to colonise harsh environments such as serum.

To determine if PdhD V76I diaphorase activity is biologically relevant we screened the impact of this variant on the NADH: NAD^+^ ratio of *S. aureus* when cultured in TSB and TSB-Serum. In both media, *pdhD* mutation leads to a decrease in the NADH: NAD+ ratio indicative of a defective TCA cycle (Figure 5A-B). The main source of cellular NADH is the TCA cycle, producing six molecules per glucose subunit. Loss of *pdhD* inhibits the pyruvate dehydrogenase complex preventing formation of acetyl-CoA, starving the TCA cycle of its main carbon source (20,30). As a result of this metabolic dysfunction, the total pool of NADH/NAD+ is reduced in *pdhD*::tn, with a 42.5% reduction when compared to JE2 in TSB (Figure S4). Complementation with the native *pdhD* restores the NADH: NAD+ ratio in both TSB and TSB-Serum, whilst restoration of the *pdhD* SNP allele only partially complemented this phenotype (Figure 5A-B). This indicates that strains expressing PdhD V76I have lower levels of NADH in comparison to wild type and native complement strain. In both complemented strains the total NADH/NAD+ pool is similar to wild type, demonstrating that generation of these coenzymes is not impacted by the PdhD V76I mutation (Figure S4). To determine if this same pattern is exhibited in the clinical strains which naturally express PdhD V76I (ASARM161, ASARM190, EOE225) we compared the NADH: NAD^+^ ratio to representative CC30 strains (ASARM4, ASARM170, EOE45) which encode wild type PdhD. Consistent with the complementation data, *S. aureus* strains expressing the *pdhD* mutant allele demonstrate a lower NADH: NAD^+^ ratio when compared to wild type (Figure 5C). This phenotype is enhanced by the presence of serum with PdhD V76I expressing strains further reducing the NADH: NAD^+^ ratio when compared to TSB only (Figure 5C).

**Figure 5.**
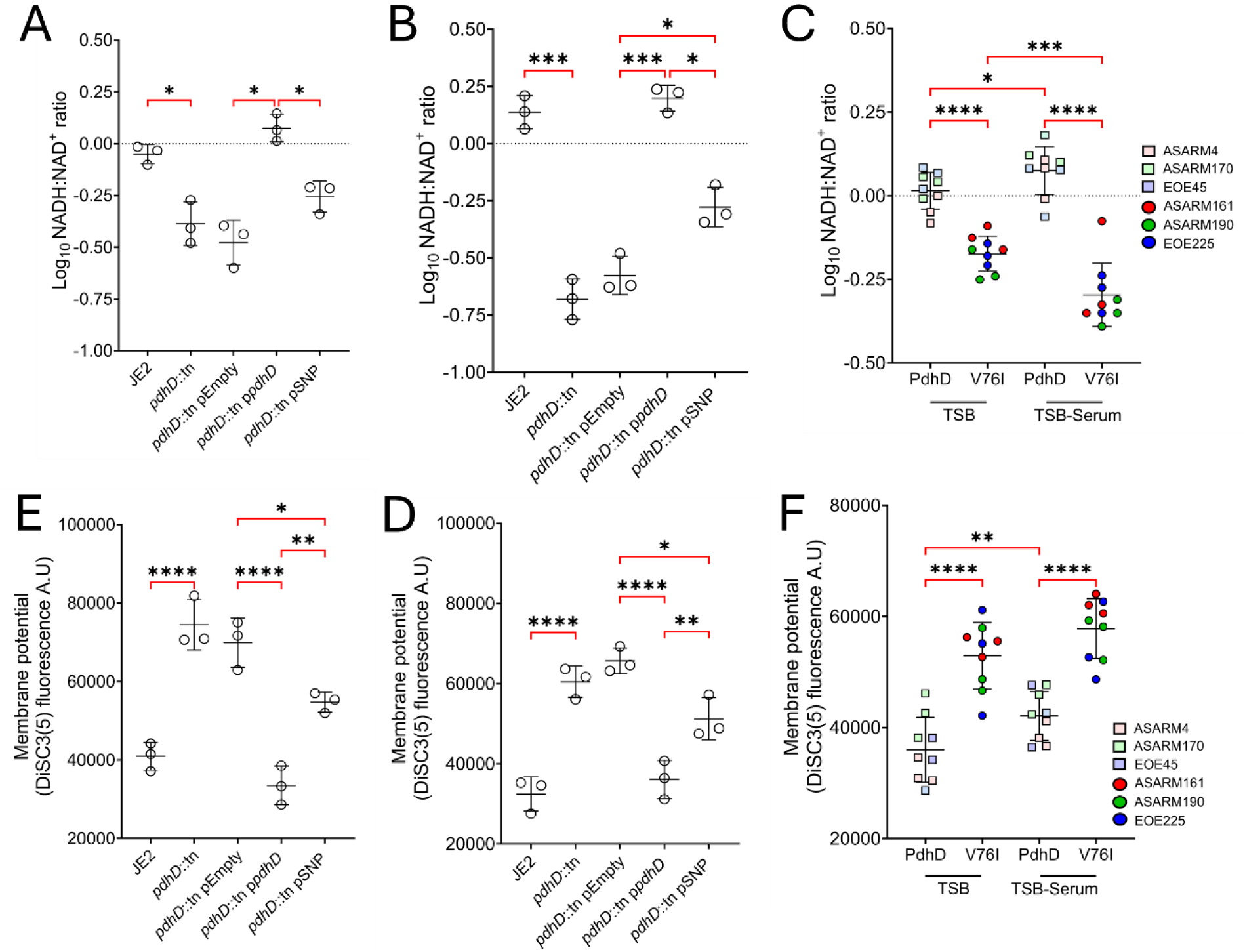
Expression of PdhD V76I is associated with reduced NADH accumulation and lower membrane potential. NADH: NAD^+^ ratio determination of isogenic *pdhD* mutants and complements in TSB (A) and TSB-Serum (B). NADH: NAD^+^ ratio determination of clinical CC30 strains containing wild type and mutant alleles of *pdhD* (C). Measurement of S. aureus membrane potential in TSB (D) and TSB-Serum (E). Membrane potential of CC30 clinical strains expressing PdhD V76I in TSB and TSB-Serum (**F**). Points represent individual experiments averaged from two technical replicates. Lines indicate the mean of these experiments with error bars denoting standard deviation. Statistical significance determined as *<0.05, **<0.01, ***<0.001, and ****<0.0001 by one-way ANOVA with Dunnett’s T3 post-hoc test (A, B, D and E) and two-way ANOVA with Tukey’s multiple comparisons test (C and F)

Lower NADH levels can be an indicator of reduced metabolic activity (20,31). NADH functions as an electron donor to the electron transport chain, serving as a substrate for type II NADH dehydrogenases. NADH dehydrogenases oxidise NADH to NAD^+^, donating 2 electrons to menaquinone (MK) to form menaquinol (MKH_2_), which is used as an electron donor for cytochrome oxidases to drive proton motive force generation (30). Therefor lower NADH levels result in decreased membrane potential, reducing the ability of *S. aureus* to produce energy and redox cycle between NADH/NAD^+^. To confirm that membrane potential is altered in PdhD V76I expressing strains we employed the voltage sensitive dye DiSC_3_(5) in both TSB and TSB-Serum. Bacteria with high negative membrane potential take up DiSC_3_(5) resulting in dye quenching, whilst strains with depolarised membranes release the dye causing an increase in fluorescence (32). As expected, loss of *pdhD* resulted in increased fluorescence in both TSB and TSB-Serum, indicative of reduced electrogenic potential and decreased metabolism (Figure 5D-E). Complementation with native *pdhD* restores membrane polarisation to wild type levels whilst strains expressing PdhD V76I also exhibit partially reduced membrane potentials, albeit not to the same extent as a *pdhD* mutant (Figure 5D-E).

We next tested the clinical strains and observed the same trend, with strains which express PdhD V76I exhibiting reduced membrane potential in comparison to representative CC30 strains (Figure 5F). This explains increased resistance to gentamicin and hydrogen peroxide of isogenic and clinical strains expressing PdhD V76I, with reduced membrane potential associated with increased resistance to both antimicrobials (27,28). Reductions in membrane potential are also associated with decreased toxin expression through repression of the Agr quorum-sensing system (33,34). Loss of *pdhD* reduced haemolysis levels to that of an *agrA* mutant indicating shut off *S. aureus* toxin production. Restoration of native *pdhD* restores haemolysis to wild type levels whilst complementation with the V76I mutant only partially compensates for loss of *pdhD,* displaying reduced haemolysis in comparison to wild type (Figure S5). This highlights how the serum resistance adaptation of strains with modified dihydrolipoamide dehydrogenase function comes at the expense of virulence potential.

### PdhD V76I diaphorase activity reduces proton flux through NADH dehydrogenases, reprogramming *S. aureus* metabolism

Whilst PdhD V76I demonstrates a reduction in DLD activity *in vitro* (Table 1) this effect is small and does not explain the eduction in membrane potential. NADH is used by the type II NADH dehydrogenases to reduce membrane bound menaquinone. If PdhD is reducing NADH more efficiently, less will cycle through NADH dehydrogenases decreasing *S. aureus* membrane potential and reducing the menaquinol: menaquinone ratio (MKH_2_: MK). To determine if this is true, we compared the MKH_2_: MK and NADH: NAD^+^ ratio of our *pdhD* mutant and complemented strains, to that of the NADH dehydrogenase mutant *ndhC*::tn. As previously reported loss of *ndhC* leads to a lower MKH_2_: MK ratios and a higher NADH: NAD^+^ ratio as the protons from NADH cannot be used to reduce MK in the bacterial membrane leading to NADH accumulation (Figure 6A-B) (30). Complementation with *ndhC* under control of its native promoter restored the ratios of both redox pairs (Figure 6A-B). Loss of *pdhD* also reduced the MKH_2_: MK and NADH: NAD^+^ ratio with reduced TCA cycle and ETC activity inhibiting production of NADH, lowering MK reduction rates (Figure 6A-B). Restoration of the wild type *pdhD* allele restored the MKH_2_: MK ratio and NADH:NAD^+^ ratio to that of JE2. Complementation with the *pdhD* SNP allele did not restore the MKH_2_: MK ratio, whilst the NADH: NAD^+^ ratio was drastically reduced as previously seen (Figure 6A-B). In terms of the MKH_2_: MK ratio, strains lacking *ndhC* and expressing PdhD V76I demonstrate similar defects in MK reduction (Figure 6A). This reduction in the MKH_2_: MK for strains expressing PdhD V76I suggests less NADH is being re-oxidised by NdhC for menaquinone reduction, with PdhD V76I contributing to NADH oxidation in the cytoplasm.

**Figure 6.**
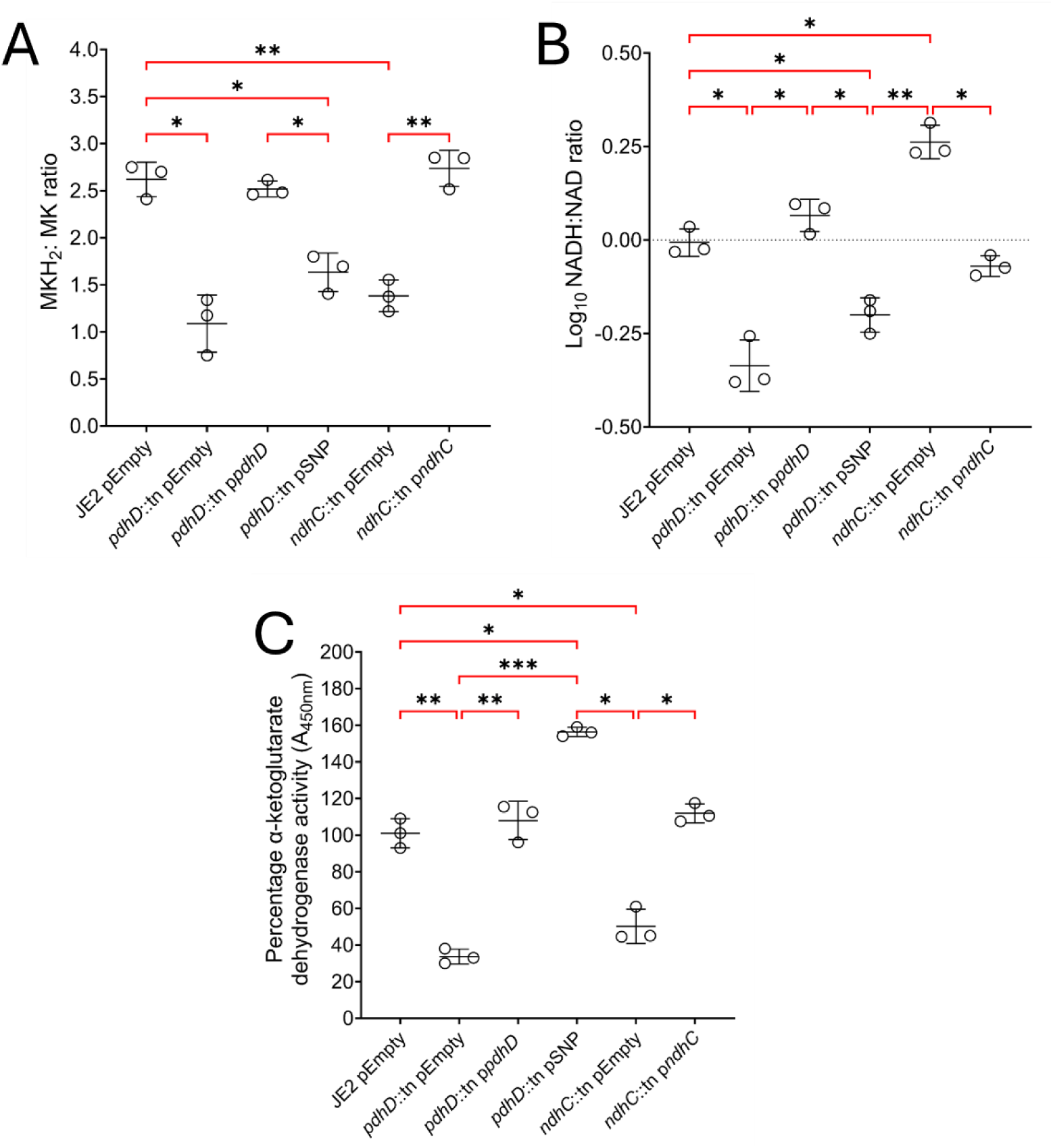
Metabolic reprogramming by PdhD V76I variant simultaneously reduces the levels of NADH and MKH_2_, impairing oxidative phosphorylation and promoting substrate-level phosphorylation. (A) Ratio of MKH_2_:MK and (B)NADH: NAD^+^ (B) in *pdhD* and *ndhC* mutants plus complements. (C) α-ketoglutarate dehydrogenase activity as a measure of TCA cycle function. Points represent individual experiments averaged from two technical replicates. Lines indicate the mean of these experiments with error bars denoting standard deviation. Significance determined as *<0.05, **<0.01, and ***<0.001, calculated by one-way ANOVA with Dunnett’s T3 multiple comparisons test.

Reduced NADH flux through NADH dehydrogenases explains the intermediate resistance of PdhD V76I expressing *S. aureus* strains to gentamicin and hydrogen peroxide, with loss of *ndhC* also demonstrating resistance to these antimicrobials due to decreased PMF (Figure S6). However, NADH dehydrogenase mutants exhibit mild growth defects in media whilst strains expressing PdhD V76I can match wild-type growth rates in liquid media. The major metabolic difference between a *ndhC*::tn mutant and strains encoding *pdhD* V76I allele is the NADH: NAD^+^ balance. One of the major products of the TCA cycle is NADH, with accumulation of NADH inhibiting TCA enzymes such as isocitrate and α-ketoglutarate dehydrogenase (35). With the level of NADH low in strains expressing PdhD V76I TCA cycle activity would not be inhibited, enabling substrate-level phosphorylation to continue when oxidative phosphorylation (respiration) is reduced. To validate this, we assessed TCA cycle activity through quantification of α-ketoglutarate dehydrogenase activity in cytoplasmic preparations derived from our isogenic *pdhD/ndhC* mutants and complements. As expected, mutation of *pdhD* and *ndhC* resulted in reduced α-ketoglutarate dehydrogenase activity, with loss of *pdhD* downregulating the TCA cycle and deletion of *ndhC* leading to accumulation of NADH (Figure 6D). Complementation with *pdhD* and *ndhC* wild-type alleles restored activity, whilst restoration of *pdhD* SNP allele increased α-ketoglutarate dehydrogenase activity by 58% in comparison to wild type and strains which are proficient for *ndhC* (Figure 6D). This confirms that whilst oxidative phosphorylation is moderately impaired in strains expressing PdhD V76I, enhanced substrate level phosphorylation (TCA cycle and glycolysis) can continue through increased redox cycling of NADH/NAD^+^.

Growth defect exhibited by *ndhC* mutants is due to shutdown of the ETC leading to accumulation of NADH, which inhibits the TCA cycle (30). To determine if the *pdhD* V76I variant can restore growth of *ndhC* mutants, the wild type and mutant *pdhD* complementation plasmids were transformed into *ndhC*::tn and growth kinetics screened in TSB. Wild type *pdhD* overexpression does not result in restoration of growth, whilst expression of PdhD V76I variant partially reduces the growth defect associated with *ndhC*::tn (Figure 7A). After 16H of culture pellets were extracted and the NADH: NAD^+^ ratio, and α-ketoglutarate dehydrogenase activity assessed. Previously loss of *ndhC* led to increased NADH accumulation (Figure 6B). Expression of the PdhD V76I variant reduced NADH accumulation in the *ndhC* mutant background leading to increased α-ketoglutarate dehydrogenase activity (Figure 7B-C). This enables partially restoration of growth in *ndhC* mutants as *S. aureus* can generate more energy through glycolysis and the TCA cycle (Figure 7A). As a result, complementation of *pdhD* and *ndhC* mutants with the PdhD V76I variant facilitates wild type growth in lab media as increased TCA cycle activity and glycolysis can counteract the defect in ETC activity. However, since normal ETC function is interrupted through loss of NADH flux through NADH dehydrogenases, increased resistance to host-derived antimicrobials such as hydrogen peroxide, AMPs and HDFAs enables strains expressing PdhD V76I to initiate active growth in serum faster than normal (Figure 1C).

**Figure 7.**
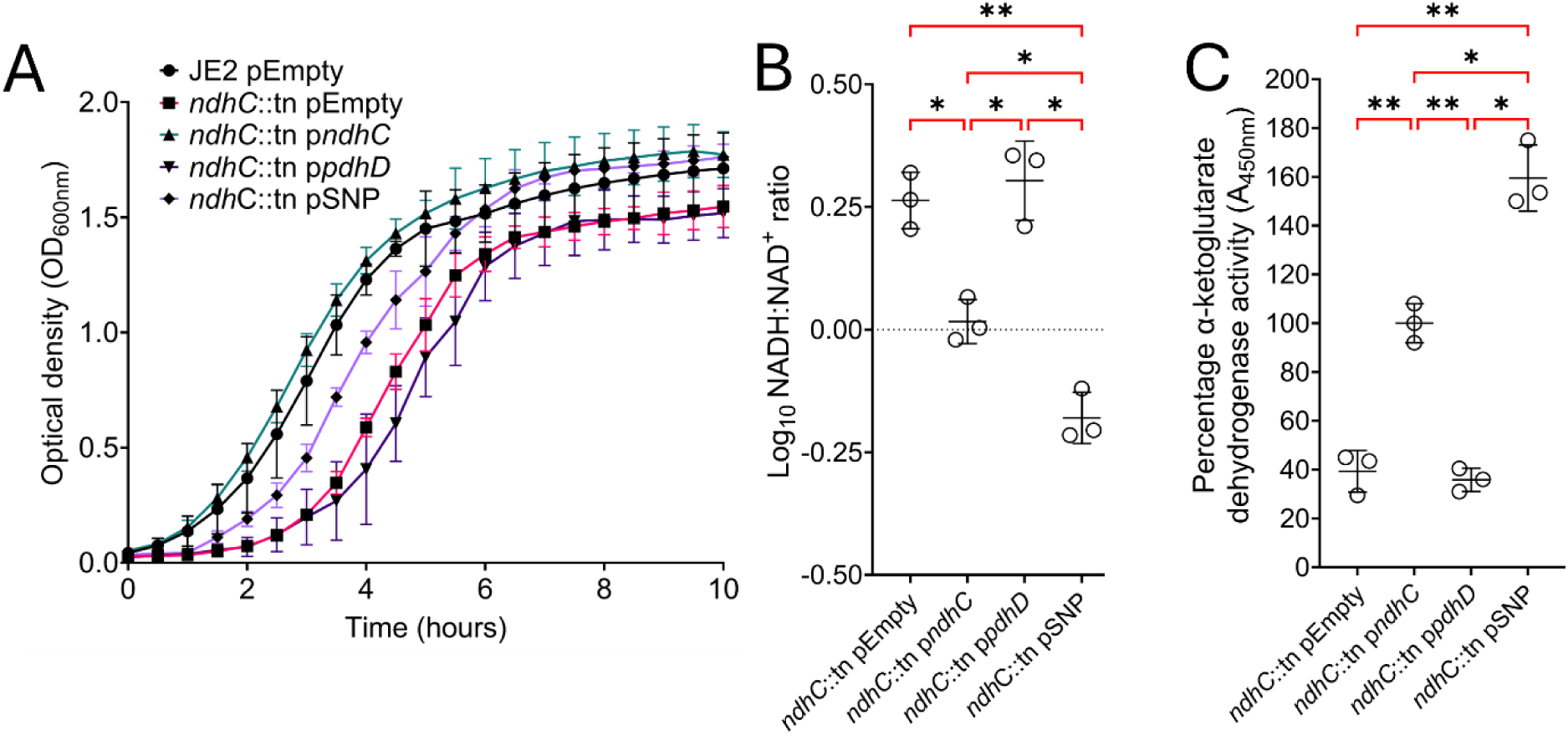
The *pdhD* SNP allele restores growth of a *ndhC* mutant through enhancing TCA cycle activity. (A) Expression of PdhD V76I in *ndhC* mutant background partially restores growth defect. (B) PdhD V76I expression in *ndhC* mutant reduces the NADH: NAD^+^ ratio, (C) and increases α-ketoglutarate dehydrogenase activity. All experiments conducted in TSB only. For A points represent the average of three experiments, error bars denote standard deviation. Optical density kinetic reads taken using a TECAN infinite pro 200 plate reader. For B and C Points represent individual experiments averaged from two technical replicates. Lines indicate the mean of these experiments with error bars denoting standard deviation. Significance determined as *<0.05, **<0.01, and ***<0.001, calculated by one-way ANOVA with Dunnett’s T3 multiple comparisons test.

## DISCUSSION

Improved survival in the bloodstream enhances the ability of *S. aureus* to sustain an infection and facilitate spread to new host sites (6). Metabolic adaptation underpins this process, with reduction in metabolic activity employed to enable *S. aureus* to survive and adapt to this hostile environment (8,17,20). As part of a previous study in which a population-based approach was employed to identify genes associated with human serum survival, a nonsynonymous amino acid substitution was identified in a dihydrolipoamide dehydrogenase (*pdhD*/*lpdA*) which enhanced fitness of *S. aureus* in human serum (23). We demonstrate that this mutation enables *S. aureus* to quickly initiate active growth in serum through alterations in NAD^+^/NADH and MK/MKH_2_ cofactor metabolism, whilst loss of *pdhD* drastically reduced survival of *S. aureus* in human serum.

Figure 8 presents a model for the metabolic scenarios in *S. aureus* strains with wild type pdhD (A), deletion of *pdhD* (B), and encoding the *pdhD* SNP allele. In strains with a normally functioning pyruvate dehydrogenase complex, pyruvate is converted into acetyl-CoA. Dihydrolipoamide dehydrogenase regenerates lipoamide from dihydrolipoamide on the E2 subunit of the pyruvate dehydrogenase, enabling the complex to continuously convert pyruvate to acetyl-CoA for the TCA cycle and fatty acid synthesis. The TCA cycle produces 6 NADH, 2 ATP, and 2 FADH_2_ per single glucose molecule, with the NADH produced subsequently utilised as a substrate by type II NADH dehydrogenases. The electrons from this oxidation reaction are used to reduce menaquinone to menaquinol. This electron donor is the used by cytochrome oxidases to drive generation of proton motive force with ATP synthase producing ATP from this proton accumulation (Figure 8A). This represents a redox-balanced energy generation system, with NADH dehydrogenase produced NAD^+^ circulating back into the TCA cycle for further NADH generation. In this scenario, *S. aureus* relies upon oxidative phosphorylation (respiration) for ATP generation yielding 30-32 ATP per glucose molecule. With complete loss of *pdhD S. aureus* metabolism shuts down (Figure 8B) (30,35,36). In pyruvate dehydrogenase deficient mutants’ acetyl-CoA cannot enter the TCA cycle. This starves the TCA cycle of its substrates, with a lack of NADH production reducing the ability of *S. aureus* to generate proton motive force and energy through oxidative phosphorylation. As a result, *S. aureus* relies upon glycolysis and acid fermentation to produce small amounts of energy and redox cycle NAD^+^/NADH coenzymes causing the reduced growth rates associated with loss of *pdhD* (Figure 8B) (20,24,34).

**Figure 8.**
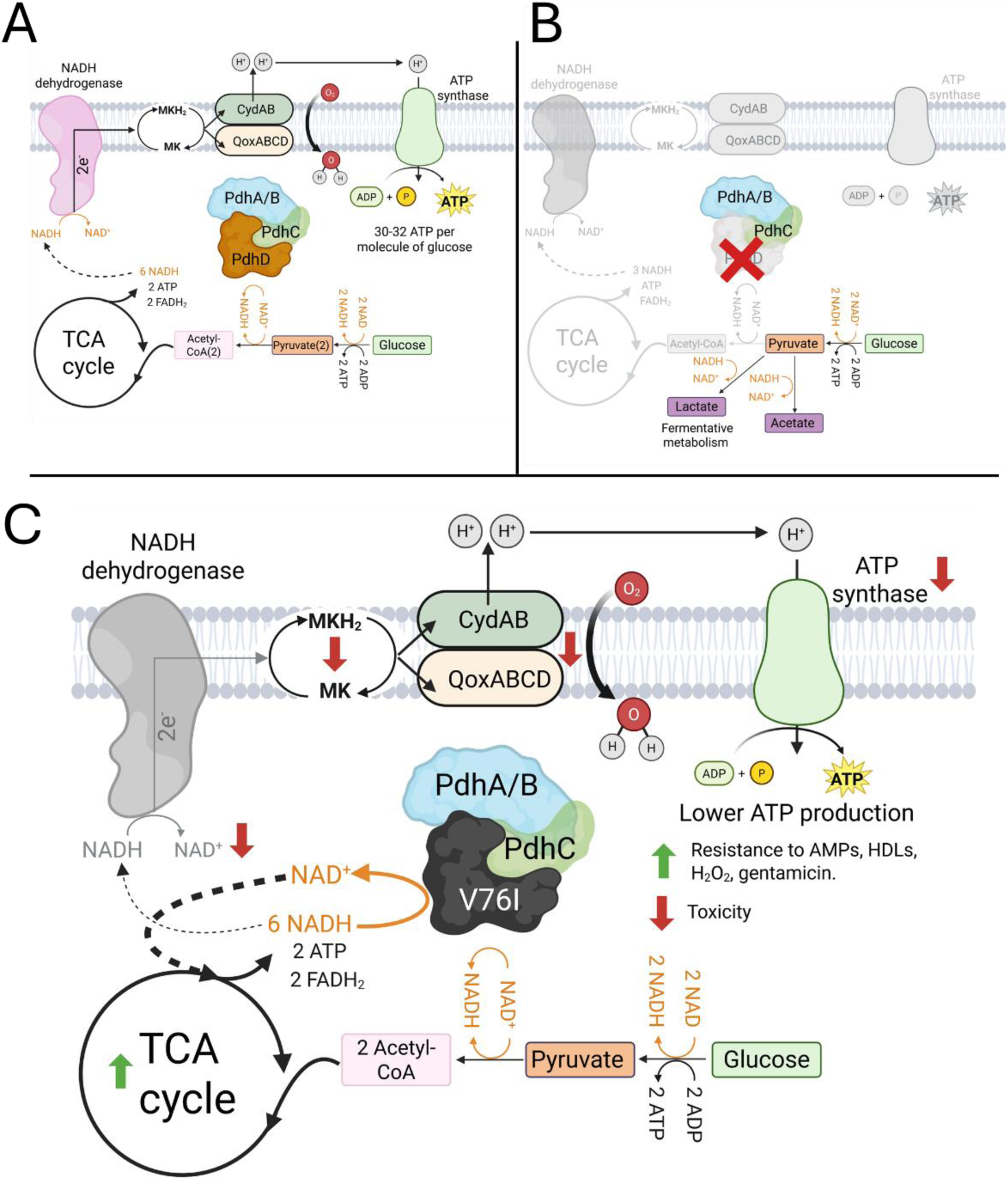
Schematic detailing metabolic outcomes for wild type (A), *pdhD*::tn (B) and *pdhD* V76I (C). (A) Strains with a functional pyruvate dehydrogenase complex generate energy and redox cycle NAD^+^/NADH through the electron transport chain and oxidative phosphorylation. Glucose is converted into 2 acetyl-CoA molecules which enter the TCA cycle, resulting in production of 6 NADH molecules. NADH dehydrogenase oxidises this coenzyme back to NAD^+^ whilst reducing menaquinone, with this electron carrier being used by cytochrome oxidases for generation of a proton gradient and subsequent ATP production. (B) Strains lacking a functional pyruvate dehydrogenase complex are unable to synthesize acetyl-CoA, starving the TCA cycle of its major carbon source. As a result, less NADH is produced inhibiting NADH dehydrogenase-dependent respiration and reducing membrane potential. Pyruvate is fermented to lactic or acetic acid for generation of small amounts of ATP and encourage limited redox cycling of NAD^+^/NADH. (C) Strains expressing the mutant PdhD V76I variant display enhanced diaphorase activity, promoting redox cycling of NADH to NAD^+^. Reduction of the intracellular NADH pool limits flux through NADH dehydrogenases, reducing efficiency of the ETC. As a result, strains expressing the PdhD V76I mutant demonstrate enhanced resistance to antimicrobials. Loss of full ETC function is compensated for through increased TCA cycle activity, and increased abundance of NAD^+^ encouraging glycolysis. Figures designed with Biorender with publication licenses g8aqbql (A), ziuf2yg (B), and oou84jt (C).

The *pdhD* SNP associated with increased survival in serum reprogrammes *S. aureus* metabolism. This V76I mutation increases the diaphorase activity of this enzyme (Figure 4D-E), allowing PdhD to contribute to redox cycling of NAD^+^/NADH, Figure 5A-C and 8C). Consequently, a partial reduction in ETC activity is observed as less NADH is available for NADH dehydrogenases to facilitate reduction of menaquinone (Figure 5D-F (30)). This increases *S. aureus* resistance to AMPs, HDLs, hydrogen peroxide and gentamicin, all of which rely upon a functional ETC to promote maximum damage (27,28,37). This resistance profile explains the ability of strains encoding the *pdhD* SNP to better tolerate serum, with AMPs and HDLs a major growth inhibition factor in this environment. Despite phenocopying *pdhD* mutant strains in exhibiting increased resistance to gentamicin, and reduced NADH, MKH_2_ and toxicity, the pathways taken to reach these phenotypes are different. Loss of *pdhD* results in metabolic shutdown, whilst strains encoding the *pdhD* SNP have redirected metabolism to favour substrate-level phosphorylation as opposed to oxidative phosphorylation (Figure 8B-C). This encourages ATP generation through glycolysis and the TCA cycle. The ability of the PdhD V76I variant to reduce NADH means fermentative metabolism of pyruvate to lactic and acetic acid are not required to recycle NADH to NAD^+^ (Figure 8B) (38). This prevents acidification of the media enabling normal cellular processes to continue and the TCA cycle to continue functioning. The lack of NADH accumulation increases TCA cycle activity, as NADH is a competitive inhibitor of α-ketoglutarate and isocitrate dehydrogenase, whilst the increased availability of NAD^+^ enhances glycolysis with this coenzyme required for glycolytic ATP generation (Figure 8 (35,39)). Consequently, strains carrying the pdhD SNP can maintain faster growth rates despite reduced oxidative phosphorylation, whilst sustaining increased levels of resistance to antimicrobials normally associated with respiratory chain defects (20,28,37). This ability to restore growth is displayed when expressing PdhD V76I in both *pdhD* and *ndhC* mutant backgrounds and represents a novel strategy of host environment adaptation (Figure 7A). Such mutations likely represent a within host evolutionary dead-end due to associated reductions in toxicity which could inhibit colonisation of both new host sites and other patients (40,41). This is especially true of bloodstream infections where the host is either killed or cured. The ability of *S. aureus* to colonise, establish and initially cause an infection is underpinned by its extensive repertoire of virulence factors. Downregulation or loss of toxicity is a common adaptation employed by *S. aureus* to promote survival *in vivo*, but entry to another site or host transmission will require these systems to be reactivated (2,40,41). Therefor whilst this mutation promotes survival during an infection, the loss of toxicity suggests that these strains would struggle to establish a new infection.

Whilst the dihydrolipoamide dehydrogenase function of this variant exhibits a small decrease in activity, it is still able to catalyse conversion from pyruvate to acetyl-CoA. This in part explains its distinct behaviour when compared to a strain devoid of *pdhD*, as enough acetyl-CoA can be generated for TCA cycle activity and fatty acid synthesis. Loss of pyruvate dehydrogenase activity results in increased membrane fluidity as straight chain fatty acids cannot be synthesized *de novo*. As a result, *S. aureus* membrane consists almost entirely of branched-chain fatty acids providing easier access to membrane targeting antimicrobials and increasing sensitivity to AMPs, HDFAs and daptomycin (24). The membrane fluidity of strains encoding the *pdhD* SNP is not obviously affected under the conditions tested in this work. However, we cannot rule out that the exact composition of fatty acids may have changed in the membrane or may be further altered under different physiological conditions.

Exactly how this PdhD alteration promotes this increased diaphorase activity is unclear. The V76I mutation does not occur in the catalytic pocket or near the co-factor binding sites of PdhD. This mutation does occur along the dimerization interface of PdhD, with dihydrolipoamide dehydrogenase activity dependent on formation of a homodimer (42,43). The increased diaphorase activity could be linked to a reduced ability to form dimers, with monomeric dihydrolipoamide dehydrogenase previously shown to exhibit enhanced diaphorase activity (42,44,45). This change to monomeric dihydrolipoamide was previously demonstrated to be driven by changes in pH however, mutations which inhibit this dimerization would have a similar effect (43). Given that *in vitro* dihydrolipoamide dehydrogenase activity only exhibits a modest drop and there is no associated growth defect with the V76I, dimerization of *pdhD* is not completely inhibited. However, a small reduction in dimerization efficiency may be driving the increased diaphorase activity exhibited by strains carrying the PdhD V76I, with this single amino acid change reprogramming *S. aureus* metabolism and promoting improved serum tolerance.

Entry and survival in human blood is a significant barrier to continuance of infection by bacteria. Mutations which promote survival in this hostile environment enable *S. aureus* to adapt faster, with persistence in this environment underpinning within host spread to other environments. This work highlights how subtle changes in central carbon metabolism can lead to drastically improved outcomes in human serum. Whilst small colony variants often result in persistent and recurrent infections, the drastic reductions in metabolic output lead to reduced growth (20,27). Through modulation of PdhD function *S. aureus* can sustain growth output whilst also maintaining enhanced resistance to host-derived antimicrobials normally associated with small colony variants. Subtle alterations in metabolism enable *S. aureus* to persist and thrive in serum, facilitating sustained infection once in a host. This highlights how complete metabolic shutdown is not the only path to survival for *S. aureus* in the hostile host environment.

## MATERIALS AND METHODS

### Bacterial strains and plasmids

Bacterial strains and plasmids used in this study are listed in table 2. *S. aureus* strains were routinely maintained on TSA and cultured in TSB with pooled human serum added at 20% (v/v) where indicated. *Escherichia coli* strains were routinely maintained on LBA and cultured in LB(Miller). For protein overexpression *E. coli* strains were cultured in Terrific Broth (TB). When required, antibiotics were added at concentrations of 10 µg mL^-1^ for erythromycin and chloramphenicol, 90 µg mL^-1^ kanamycin, and 100 µg mL^-1^ carbenicillin.

**Table 2.**
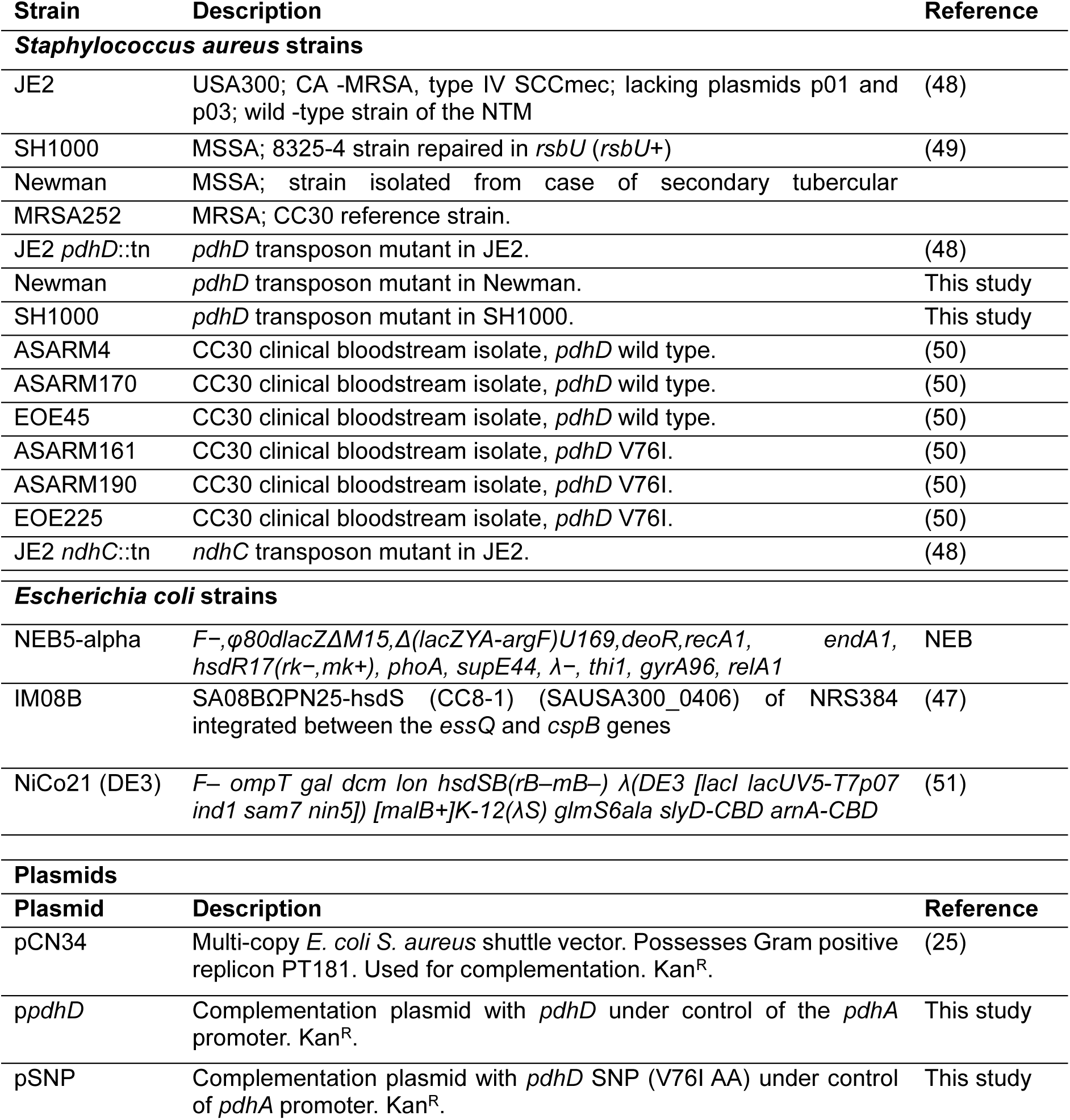
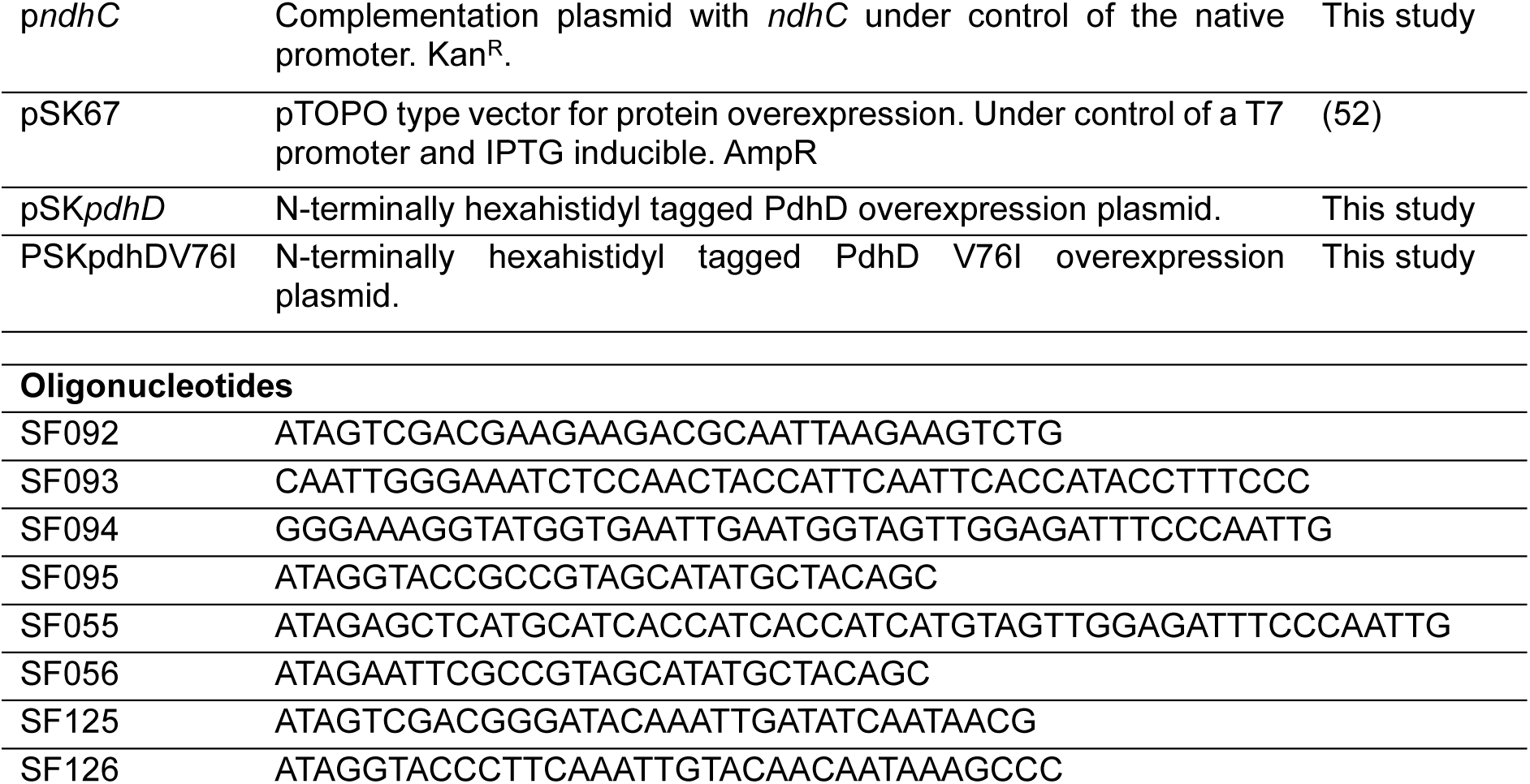
Strains, plasmids and oligonucleotides used in this study.

### DNA manipulations, mutant construction and complementation

Plasmid minipreps and PCR purifications performed with Qiaprep spin mini kit (Qiagen). PCR products for cloning applications amplified with Phusion polymerase (Thermo). Restriction digest, ligations and transformation performed according to standard protocols. Oligonucleotide sequences (Sigma) used in this study are in table 2.

The *pdhD*::tn mutation from JE2 was transduced into SH1000 using φ11 as performed by Krausz and Bose (46). As *pdhD* is the last gene in the *pdhABCD* operon we placed *pdhD* under control of the native *pdhA* promoter in our complementation plasmids. The P*_pdhA_* promoter, *pdhD* coding sequence and downstream terminator regions were amplified from MRSA252 with primers SF092/93 and SF094/95 respectively. P*_pdhA_* and *pdhD* DNA sequences were joined by overlap extension PCR through a 21bp overlapping region on SF093 and SF094. The P*_pdhA_*-*pdhD* fragment was then ligated into the *E. coli*/*S. aureus* shuttle vector pCN34 through digestion with KpnI and SalI. Ligations were transformed into *E. coli* IM08B, purified, and then electroporated into the required *S. aureus* strains (47). For generation of *pdhD* V76I complementation plasmid the same process was followed with the *pdhD* coding sequence and terminator region amplified from ASARM161.

### Serum survival assays

Pooled human serum was purchased (Clinisciences), aliquoted and stored at -20^°^C until use. *S. aureus* strains were cultured in TSB O/N, normalised to 1x10^7^ CFU mL^-1^ in PBS and 20 µL inoculated into 180 µL 25% human serum diluted in PBS. The bacteria were then incubated at 37^°^C with shaking, with three samples sacrificed per timepoint for bacterial enumeration. The same number of bacterial cells were inoculated into PBS, diluted, and plated as a control. Serial dilutions were plated on TSA to determine CFU mL^-1^. Survival was determined as the percentage CFU in serum relative to the initial inoculum (0 H timepoint). Experiments repeated three times with three technical replicates per timepoint.

### Antimicrobial killing assays

Cationic antimicrobial peptide killing assays with LL37/HNP-1 (Clinisciences) and hydrogen peroxide were performed in 1 mL of PBS with a 1x10^6^ *S. aureus* inoculum. Both LL37 and HNP-1 were added at 5 µg mL^-1^ and incubated at 37^°^C with shaking for 4 H. Hydrogen peroxide was added at 20 mM and incubated at 37^°^C with shaking for 90 minutes. 1 mL of Dey-Engley broth was added to hydrogen peroxide killing assays to neutralise and prevent further killing during sample processing. Killing assays with arachidonic acid (Sigma), gentamicin (Sigma) and daptomycin (Med-Chem express) were performed in 1 mL TSB with a starting inoculum of 1x10^7^ cells. Arachidonic acid and gentamicin were added at 200 µM and 5 µg mL^-1^ respectively, followed by incubation at 37^°^C for 2 H.. Data presented as the base Log^10^ of the raw CFU data. The same number of bacterial cells inoculated into PBS, diluted, and plated acted as a control.

### Laurdan membrane fluidity assay

Laurdan dye (MedChem) was used to assess membrane fluidity as previously described by Wenzel and colleagues (26). *S. aureus* strains were cultured overnight in TSB and sub-cultured into 5 mL TSB supplemented with 1.25 mM CaCl_2_, 0.5 mM MgCl_2_, and 0.2% glucose (w/v) the following day. The bacteria were incubated for 3 H at 37^°^C, normalised to an OD_600nm_ of 0.4 in the same media (pre-warmed to 37^°^C), and stained with 10 µM of Laurdan dye for 5 minutes. Cells were washed three times in pre-warmed PBS supplemented with 1.25 mM CaCl_2_, 0.5 mM MgCl_2_, and 0.2% glucose, with samples incubated in a dry water bath set to 37^°^C during processing. The cell suspension (200 µL) was added to opaque black 96-well plates (Costar) and membrane fluidity determined by measuring fluorescence (excitation 330nm; emission 460 and 500nm) using a TECAN PRO200 Infinite plate reader. Generalised polarisation value calculated using formula: GP = (I_460_-I_500_)/(I_460_+I_500_) where I_460_ and I_500_ are the emission intensities at 460 and 500nm respectively. DMSO (1%) and benzyl alcohol (1%) added to JE2 as negative and positive controls respectively.

### PdhD overexpression and purification

The *pdhD* gene was amplified from MRSA252 (wild type) and ASARM161 (V76I) using primer pair SF055/56 modified with a hexahistidyl tag and EcoRI/SacI restriction sites. The modified *pdhD* DNA fragments were cloned into the IPTG-inducible plasmid pSK67 using the EcoRI/SacI restriction sites, forming plasmids pSK*pdhD* and pSK*pdhD*V76I. Plasmid insert sequences were confirmed by Sanger sequencing (Eurofins). Vectors were transformed into *E. coli* NiCo21 (DE3) and selected for with carbenicillin. Single colonies were selected and cultured in LB broth for 18 H at 37°C. Strains were then sub-cultured to an OD600nm of 0.05 in TB and grown to an OD of 0.6-0.8. Cultures were then cooled to 20°C for 45 minutes and induced with 0.2 mM isopropyl-β-d-thiogalactopyranoside. Following 18 H of culture at 20°C cell pellets were harvested by centrifugation and stored at -70°C for at least 1 day prior to processing. For purification of soluble PdhD and PdhD V76I immobilised metal ion affinity chromatography (IMAC) followed by buffer exchange with a PD-10 desalting column was used. Frozen pellets were defrosted and resuspended in lysis buffer consisting of 50 mM Tris-HCl (pH 8.0), 150 mM NaCl, 10 mM Imidazole, 1.2 µg mL^-1^ lysozyme, and a protease inhibitor cocktail (Thermo Scientfic). Pellets were resuspended at 1g of cell paste per 10 mL of lysis buffer. Cells were lysed with a sonic dismembranator with a 15s/15s ON/OFF cycle, samples were stored in ice throughout the sonication process.

IMAC was performed using a HiTrap Chelating HP column (Cytiva) charged with NiCl_2_. The column was equilibrated with 10 column volumes (CV) of buffer A containing 50 mM Tris-HCl (pH 8.0), 500 mM NaCl, 10 mM Imidazole, and 5% (v/v) glycerol with a P1 peristaltic pump (Cytiva). Samples containing PdhD and PdhD V76I were applied to the equilibrated column at 1 mL min^-1^. Soluble PdhD and PdhD V76I were obtained with a stepwise imidazole gradient with 5 CV washes at concentrations of 10-, 20-, 40- and 80-mM imidazole. Samples were then eluted with 3 CV of Buffer A containing 200- and 500-mM imidazole collected at 2.5 mL aliquots. PdhD samples were then analysed by sodium dodecyl sulphate-polyacrylamide gel electrophoresis (SDS-PAGE) for homogeneity. Pure samples were pooled and concentrated using a NeoSpin centrifugal concentrator (10kDa cutoff). Concentrated PdhD were then applied to PD-10 desalting columns (Cytiva) and eluted with 20 mM Tris-HCl (pH 8.0), 150 mM NaCl and reconcentrated to 10 mg mL^-1^. Samples were aliquoted, flash frozen in liquid nitrogen, and stored at -70 for use as required. Prior to freezing 5% glycerol was added as a cryoprotectant. After defrosting, samples were resuspended to 2.5 mL in 50 mM Tris-HCl (pH 8.0), 150 mM NaCl and glycerol removed by buffer exchange with a PD-10 column.

### Dihydrolipoamide dehydrogenase enzyme assay

DLD activity was assayed as described previously (53) with minor modifications. The reactions were performed in 96-well plates with a final reaction volume of 250 µL. The initial reaction mixture contained 100 mM Potassium phosphate buffer (pH 7.8), 1 mM EDTA, 0.4 mM DHL, 0.3 mM NAD^+^, and 0.003 mM of PdhD. Reactions started upon addition of DHL and monitored through NADH production at A_340nm_. Reactions containing all components except DHL used as blanks. For kinetic parameter quantification variable concentrations of DHL and NAD^+^ were used between 0-20 mM. When DHL was varied the reaction was saturated with 20mM NAD^+^. When NAD^+^ was varied the reaction was saturated with 20 mM DHL. Kinetic constants (V_max_, K_m_ and K_cat_) determined using non-linear regression in GraphPad prism (version 10.4.1)

### Diaphorase enzyme activity

DP activity of PdhD variants measured using thiazolyl blue tetrazolium bromide (MTT) as done previously (43,45). The reactions were performed in clear flatbottomed 96-well plates with a final reaction volume of 250 µL. The initial reaction mixture contained 100 mM Potassium phosphate buffer (pH 7.6), 0.4 mM MTT, 0.3 mM NADH, and 0.003 mM of PdhD. Reactions started upon addition of MTT and monitored through production of the blue-purple coloured formazan following reduction of MTT at A_560nm_ using a TECAN infinite 200 plate reader. For kinetic parameter quantification variable concentrations of MTT and NADH were used between 0-10 mM. When MTT was varied the reaction was saturated with 10 mM NADH. When NADH was varied the reaction was saturated with 10 mM MTT. Kinetic constants (V_max_, K_m_ and K_cat_) determined using non-linear regression in GraphPad prism (version 10.4.1)

### NADH: NAD^+^ ratio determination

NADH: NAD^+^ ratio quantified using a colorimetric assay kit (Novus) according to manufacturer’s instructions. Samples prepared by mechanically lysing 1X10^8^ *S. aureus* cells after 16 H of culture in 3 mL of TSB or TSB with 20% human serum. Cells were resuspended in 500 µL assay buffer, and 0.1 g of glass beads (Sigma) added to sample tubes. Mechanical lysis performed at 6 M/s for 20s repeated 3 times using a

MP fast prep tissue homogeniser with samples stored on ice throughout. Prior to NAD^+^/NADH quantification samples were passed through a 3kDa centrifugal concentrator to remove protein contaminants. Colorimetric change measured in clear flatbottomed 96-well plates with a TECAN infinite 200 plate reader. Figures represented as the base log_10_ of the calculated NADH: NAD^+^ ratios.

### Membrane polarity measurement

Membrane polarity was measured using 3,3′-dipropylthiadicarbocyanine iodide (DiSC_3_(5)), Thermofisher Scientific) as described previously (54) with minor modifications. Samples cultured in 3 mL TSB or TSB with 20% human serum diluted to an OD_600nm_ of 0.4 in prewarmed (37^°^C) TSB. To black flat bottomed 96-well plates 200 µL of culture added per well, and DiSC_3_(5) added to a final concentration of 1 µM. Cultures were mixed with dye and then incubated statically at 37°C for 5 minutes. Fluorescence was then measured using a TECAN infinite 200 plate reader (excitation 622nm; emission 670nm). Fluorescence values divided by OD_600nm_ measurements to normalise to each samples cell density.

### XTT metabolic reduction assay

XTT reduction assay performed according to Xu et al (55) with minor changes. *S. aureus* were cultured overnight in TSB at 37°C. Strains were diluted 1:50 and reinoculated into fresh TSB followed by further incubation at 37°C for 3 H. Cells were diluted to 1X10^8^ CFU mL^-1^ in PBS, with 100 µL added to a flat-bottomed 96 well plate. Immediately 100 µL of a XTT (1mg/mL Sigma) and menadione (0.4 mM Sigma) mixture was added to each well and incubated at 37°C for 3 H. XTT reduction was measured in a TECAN PRO200 infinite plate reader at A_490nm_. A well with sterile media and the XTT-menadione solution were used as a blank.

### Alpha-ketoglutarate dehydrogenase activity assay

Alpha-ketoglutarate dehydrogenase activity assessed using a colorimetric assay kit (Novus) according to manufacturer’s instructions. Samples prepared by mechanically lysing 1x10^8^ *S. aureus* cells after 16 H incubation in 3 mL TSB. Mechanical lysis procedure identical to that for NADH: NAD^+^ ratio determination (above). Prior to activity assessment, lysate was filtered was filtered through a 10 kDa centrifugal concentrator to remove any excess α-ketoglutarate and pyruvate (interferes with assay). Protein containing samples were resuspended in 20 mM Tris-HCl (pH 7.4), 150 mM NaCl.

## ACKNOWLEDGMENTS

All authors acknowledge the provision of strain by the Network on Antimicrobial Resistance in *Staphylococcus aureus* (NARSA) Program: under NIAID/NIH Contract number HHSN272200700055C. This work was supported by Research Ireland Frontier for the Future Program Award (reference: 21/FFP-A/9704), and a Wellcome Trust Investigator Award (reference: 212258/Z/18/Z).

## DATA AVAILIBILITY STATEMENT

The authors confirm that the data supporting the findings of this study are available with the article and supplementary material. Bacterial strains developed as part of this study are available upon request.

## REFERENCES

1. Antimicrobial Resistance Collaborators. Global burden of bacterial antimicrobial resistance in 2019: a systematic analysis. Lancet Lond Engl. 2022 Feb 12;399(10325):629–55.

2. Gordon RJ, Lowy FD. Pathogenesis of methicillin-resistant Staphylococcus aureus infection. Clin Infect Dis Off Publ Infect Dis Soc Am. 2008 June 1;46 Suppl 5(Suppl 5):S350–359.

3. Kaasch AJ, Barlow G, Edgeworth JD, Fowler VG, Hellmich M, Hopkins S, et al. Staphylococcus aureus bloodstream infection: a pooled analysis of five prospective, observational studies. J Infect. 2014 Mar;68(3):242–51.

4. Bai AD, Lo CKL, Komorowski AS, Suresh M, Guo K, Garg A, et al. Staphylococcus aureus bacteraemia mortality: a systematic review and meta-analysis. Clin Microbiol Infect Off Publ Eur Soc Clin Microbiol Infect Dis. 2022 Aug;28(8):1076–84.

5. Tong SYC, Fowler VG, Skalla L, Holland TL. Management of Staphylococcus aureus Bacteremia: A Review. JAMA. 2025 Sept 2;334(9):798–808.

6. Kwiecinski JM, Horswill AR. Staphylococcus aureus bloodstream infections: pathogenesis and regulatory mechanisms. Curr Opin Microbiol. 2020 Feb;53:51–60.

7. Kahl BC, Becker K, Löffler B. Clinical Significance and Pathogenesis of Staphylococcal Small Colony Variants in Persistent Infections. Clin Microbiol Rev. 2016 Apr;29(2):401–27.

8. Rowe SE, Beam JE, Conlon BP. Recalcitrant Staphylococcus aureus Infections: Obstacles and Solutions. Infect Immun. 2021 Mar 17;89(4):e00694–20.

9. Grosz M, Kolter J, Paprotka K, Winkler AC, Schäfer D, Chatterjee SS, et al. Cytoplasmic replication of Staphylococcus aureus upon phagosomal escape triggered by phenol-soluble modulin α. Cell Microbiol. 2014 Apr;16(4):451–65.

10. Tam K, Torres VJ. Staphylococcus aureus Secreted Toxins and Extracellular Enzymes. Microbiol Spectr. 2019 Mar;7(2).

11. Laabei M, Uhlemann AC, Lowy FD, Austin ED, Yokoyama M, Ouadi K, et al. Evolutionary Trade-Offs Underlie the Multi-faceted Virulence of Staphylococcus aureus. PLoS Biol. 2015;13(9):e1002229.

12. Rose HR, Holzman RS, Altman DR, Smyth DS, Wasserman GA, Kafer JM, et al. Cytotoxic Virulence Predicts Mortality in Nosocomial Pneumonia Due to Methicillin-Resistant Staphylococcus aureus. J Infect Dis. 2015 June 15;211(12):1862–74.

13. Ganesan N, Mishra B, Felix L, Mylonakis E. Antimicrobial Peptides and Small Molecules Targeting the Cell Membrane of Staphylococcus aureus. Microbiol Mol Biol Rev MMBR. 2023 June 28;87(2):e0003722.

14. Levy O. Antimicrobial proteins and peptides of blood: templates for novel antimicrobial agents. Blood. 2000 Oct 15;96(8):2664–72.

15. Das UN. Arachidonic acid and other unsaturated fatty acids and some of their metabolites function as endogenous antimicrobial molecules: A review. J Adv Res. 2018 May;11:57–66.

16. Beavers WN, Monteith AJ, Amarnath V, Mernaugh RL, Roberts LJ, Chazin WJ, et al. Arachidonic Acid Kills Staphylococcus aureus through a Lipid Peroxidation Mechanism. mBio. 2019 Oct 1;10(5):e01333–19.

17. Chang J, Lee C, Kim I, Kim J, Kim JH, Yun T, et al. Environmental cues in different host niches shape the survival fitness of Staphylococcus aureus. Nat Commun. 2025 July 28;16(1):6928.

18. Cassat JE, Skaar EP. Metal ion acquisition in Staphylococcus aureus: overcoming nutritional immunity. Semin Immunopathol. 2012 Mar;34(2):215–35.

19. Healy C, Munoz-Wolf N, Strydom J, Faherty L, Williams NC, Kenny S, et al. Nutritional immunity: the impact of metals on lung immune cells and the airway microbiome during chronic respiratory disease. Respir Res. 2021 Apr 29;22(1):133.

20. Proctor R. Respiration and Small Colony Variants of Staphylococcus aureus. Microbiol Spectr. 2019 May;7(3).

21. Gaupp R, Schlag S, Liebeke M, Lalk M, Götz F. Advantage of upregulation of succinate dehydrogenase in Staphylococcus aureus biofilms. J Bacteriol. 2010 May;192(9):2385–94.

22. Bohn C, Rigoulay C, Chabelskaya S, Sharma CM, Marchais A, Skorski P, et al. Experimental discovery of small RNAs in Staphylococcus aureus reveals a riboregulator of central metabolism. Nucleic Acids Res. 2010 Oct;38(19):6620–36.

23. Douglas EJA, Palk N, Brignoli T, Altwiley D, Boura M, Laabei M, et al. Extensive remodelling of the cell wall during the development of Staphylococcus aureus bacteraemia. eLife. 2023 July 4;12:RP87026.

24. Singh VK, Sirobhushanam S, Ring RP, Singh S, Gatto C, Wilkinson BJ. Roles of pyruvate dehydrogenase and branched-chain α-keto acid dehydrogenase in branched-chain membrane fatty acid levels and associated functions in Staphylococcus aureus. J Med Microbiol. 2018 Apr;67(4):570–8.

25. Charpentier E, Anton AI, Barry P, Alfonso B, Fang Y, Novick RP. Novel cassette-based shuttle vector system for gram-positive bacteria. Appl Environ Microbiol. 2004 Oct;70(10):6076–85.

26. Wenzel M, Vischer NOE, Strahl H, Hamoen LW. Assessing Membrane Fluidity and Visualizing Fluid Membrane Domains in Bacteria Using Fluorescent Membrane Dyes. Bio-Protoc. 2018 Oct 20;8(20):e3063.

27. Baumert N, von Eiff C, Schaaff F, Peters G, Proctor RA, Sahl HG. Physiology and antibiotic susceptibility of Staphylococcus aureus small colony variants. Microb Drug Resist Larchmt N. 2002;8(4):253–60.

28. Painter KL, Strange E, Parkhill J, Bamford KB, Armstrong-James D, Edwards AM. Staphylococcus aureus adapts to oxidative stress by producing H2O2-resistant small-colony variants via the SOS response. Infect Immun. 2015 May;83(5):1830–44.

29. Patel MS, Nemeria NS, Furey W, Jordan F. The pyruvate dehydrogenase complexes: structure-based function and regulation. J Biol Chem. 2014 June 13;289(24):16615–23.

30. Schurig-Briccio LA, Parraga Solorzano PK, Lencina AM, Radin JN, Chen GY, Sauer JD, et al. Role of respiratory NADH oxidation in the regulation of Staphylococcus aureus virulence. EMBO Rep. 2020 May 6;21(5):e45832.

31. Somerville GA, Proctor RA. At the crossroads of bacterial metabolism and virulence factor synthesis in Staphylococci. Microbiol Mol Biol Rev MMBR. 2009 June;73(2):233–48.

32. Te Winkel JD, Gray DA, Seistrup KH, Hamoen LW, Strahl H. Analysis of Antimicrobial-Triggered Membrane Depolarization Using Voltage Sensitive Dyes. Front Cell Dev Biol. 2016;4:29.

33. Douglas EJA, Duggan S, Brignoli T, Massey RC. The MpsB protein contributes to both the toxicity and immune evasion capacity of Staphylococcus aureus. Microbiol Read Engl. 2021 Oct;167(10):001096.

34. Tuchscherr L, Löffler B, Proctor RA. Persistence of Staphylococcus aureus: Multiple Metabolic Pathways Impact the Expression of Virulence Factors in Small-Colony Variants (SCVs). Front Microbiol. 2020;11:1028.

35. Maynard AG, Kanarek N. NADH Ties One-Carbon Metabolism to Cellular Respiration. Cell Metab. 2020 Apr 7;31(4):660–2.

36. Arumugam P, Kielian T. Metabolism Shapes Immune Responses to Staphylococcus aureus. J Innate Immun. 2024;16(1):12–30.

37. Gläser R, Becker K, von Eiff C, Meyer-Hoffert U, Harder J. Decreased susceptibility of Staphylococcus aureus small-colony variants toward human antimicrobial peptides. J Invest Dermatol. 2014 Sept;134(9):2347–50.

38. Troitzsch A, Loi VV, Methling K, Zühlke D, Lalk M, Riedel K, et al. Carbon Source-Dependent Reprogramming of Anaerobic Metabolism in Staphylococcus aureus. J Bacteriol. 2021 Mar 23;203(8):e00639–20.

39. Hopp AK, Grüter P, Hottiger MO. Regulation of Glucose Metabolism by NAD+ and ADP-Ribosylation. Cells. 2019 Aug 13;8(8):890.

40. Giulieri SG, Guérillot R, Duchene S, Hachani A, Daniel D, Seemann T, et al. Niche-specific genome degradation and convergent evolution shaping Staphylococcus aureus adaptation during severe infections. eLife. 2022 June 14;11:e77195.

41. Laabei M, Uhlemann AC, Lowy FD, Austin ED, Yokoyama M, Ouadi K, et al. Evolutionary Trade-Offs Underlie the Multi-faceted Virulence of Staphylococcus aureus. PLoS Biol. 2015;13(9):e1002229.

42. Tsai CS, Templeton DM, Wand AJ. Multifunctionality of lipoamide dehydrogenase: activities of chemically trapped monomeric and dimeric enzymes. Arch Biochem Biophys. 1981 Jan;206(1):77–86.

43. Klyachko NL, Shchedrina VA, Efimov AV, Kazakov SV, Gazaryan IG, Kristal BS, et al. pH-dependent substrate preference of pig heart lipoamide dehydrogenase varies with oligomeric state: response to mitochondrial matrix acidification. J Biol Chem. 2005 Apr 22;280(16):16106–14.

44. Babady NE, Pang YP, Elpeleg O, Isaya G. Cryptic proteolytic activity of dihydrolipoamide dehydrogenase. Proc Natl Acad Sci U S A. 2007 Apr 10;104(15):6158–63.

45. Vaubel RA, Rustin P, Isaya G. Mutations in the dimer interface of dihydrolipoamide dehydrogenase promote site-specific oxidative damages in yeast and human cells. J Biol Chem. 2011 Nov 18;286(46):40232–45.

46. Krausz KL, Bose JL. Bacteriophage Transduction in Staphylococcus aureus: Broth-Based Method. Methods Mol Biol Clifton NJ. 2016;1373:63–8.

47. Monk IR, Tree JJ, Howden BP, Stinear TP, Foster TJ. Complete Bypass of Restriction Systems for Major Staphylococcus aureus Lineages. mBio. 2015 May 26;6(3):e00308–00315.

48. Fey PD, Endres JL, Yajjala VK, Widhelm TJ, Boissy RJ, Bose JL, et al. A genetic resource for rapid and comprehensive phenotype screening of nonessential Staphylococcus aureus genes. mBio. 2013 Feb 12;4(1):e00537–00512.

49. Horsburgh MJ, Aish JL, White IJ, Shaw L, Lithgow JK, Foster SJ. sigmaB modulates virulence determinant expression and stress resistance: characterization of a functional rsbU strain derived from Staphylococcus aureus 8325-4. J Bacteriol. 2002 Oct;184(19):5457–67.

50. Recker M, Laabei M, Toleman MS, Reuter S, Saunderson RB, Blane B, et al. Clonal differences in Staphylococcus aureus bacteraemia-associated mortality. Nat Microbiol. 2017 Oct;2(10):1381–8.

51. Robichon C, Luo J, Causey TB, Benner JS, Samuelson JC. Engineering Escherichia coli BL21(DE3) derivative strains to minimize E. coli protein contamination after purification by immobilized metal affinity chromatography. Appl Environ Microbiol. 2011 July;77(13):4634–46.

52. Fenn S, Dubern JF, Cigana C, De Simone M, Lazenby J, Juhas M, et al. NirA Is an Alternative Nitrite Reductase from Pseudomonas aeruginosa with Potential as an Antivirulence Target. mBio. 2021 Apr 20;12(2):e00207–21.

53. Wei W, Li H, Nemeria N, Jordan F. Expression and purification of the dihydrolipoamide acetyltransferase and dihydrolipoamide dehydrogenase subunits of the Escherichia coli pyruvate dehydrogenase multienzyme complex: a mass spectrometric assay for reductive acetylation of dihydrolipoamide acetyltransferase. Protein Expr Purif. 2003 Mar;28(1):140–50.

54. Ledger EVK, Edwards AM. Growth Arrest of Staphylococcus aureus Induces Daptomycin Tolerance via Cell Wall Remodelling. mBio. 2023 Feb 28;14(1):e0355822.

55. Xu Z, Liang Y, Lin S, Chen D, Li B, Li L, et al. Crystal Violet and XTT Assays on Staphylococcus aureus Biofilm Quantification. Curr Microbiol. 2016 Oct;73(4):474–82.

