## Supplementary material for "Metabolic reprogramming promotes *Staphylococcus aureus* serum resistance during bacteraemia"

Commented [RM1]: ??

Commented [SF2R1]: Yep, much better. Reenforces the biological relevance.

**Samuel Fenn[1,2], Ruth C. Massey[1, 2, 3, *]**

[1] School of Microbiology, University College Cork, Cork, Ireland.

[2] APC Microbiome, University College Cork, Cork, Ireland.

[3] School of Cellular and Molecular Medicine, University of Bristol, Bristol, UK.

[*] Corresponding author

**SUPPLEMENTARY FIGURES AND TABLES**

**Figure S1 –** Expression of the PdhD V76I variant enhances survival and encourages growth of *S. aureus* in human serum.

**Figure S2 -** Sensitivity of clinical CC30 strains to LL37, HNP-1 and Arachidonic Acid.

**Figure S3 -** Growth kinetics of CC30 clinical strains in TSB (A) and RPMI-C (B).

**Table S1 -** Minimum inhibitory concentration assays for isogenic JE2 and SH1000 mutants with associated complements.

**Table S2** - Kinetic parameters assessing activity of PdhD vs PdhD V76I.

**Figure S4 -** Total NAD/NADH pool in JE2 mutant and complement strains in TSB (A) and TSB-Serum (B).

**Figure S5 –** PdhD V76I mutation is associated with reduced haemolysis.

**Figure S6** – Loss of *ndhC* phenocopies PdhD V76I increased resistance to gentamicin (A) and hydrogen peroxide (B).

**Figure S1 - Expression of the PdhD V76I variant enhances survival and encourages growth of *S. aureus* in human serum.** Tracking of SH1000 isogenic mutant strains and complements serum survival across an 8-hour incubation window at 37˚C. Complementation with *pdhD* restores serum survival to SH1000 to wild type levels. Restoration of *pdhD* V76I increases serum survival and encourages active growth of *S. aureus* in serum from 2 hours onwards, outperforming both SH1000 wild type and the native *pdhD* complement strain. Serum survival with SH1000 strains and serum killing over time experiments averaged from three biological replicates with a minimum of three technical replicates. Percentage values normalised to initial inoculum of strains at hour 0. Error bars represent standard deviation. Significance determined as *<0.05, **<0.01, calculated by Two Way ANOVA with Tukey's multiple comparisons test.

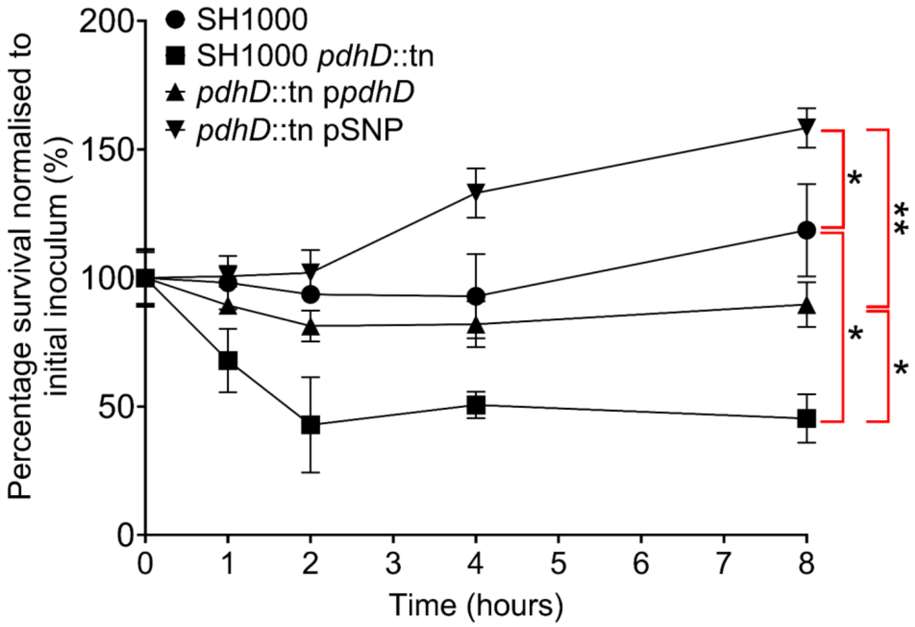

**Figure S2 – Sensitivity of clinical CC30 strains to LL37, HNP-1 and Arachidonic Acid.** Difference in susceptibility between CC30 strains with wild type PdhD (ASARM4, ASARM170 and EOE45) and PdhD V76I (ASARM161, ASARM190 and EOE225). Clinical strains expressing PdhD V76I demonstrate enhanced resistance to LL37, HNP-1 and AA in comparison to wild type PdhD. LL37 and HNP-1 killing assays conducted in PBS at 5 µg $ml^{-1}$ for 4 hours with a starting inoculum of $1x10^6$ cells. AA killing assays conducted in TSB at 200 µM and 2.5 µg $ml^{-1}$ for 2 hours with a starting inoculum of $1x10^7$ cells. Each dot represents and individual experiment with a minimum of three technical replicates technical replicates. Bars represent man values, error bars the standard deviation. Significance determined as **<0.01, ****<0.0001 calculated by One Way ANOVA with Dunnett's T3 multiple comparisons test.

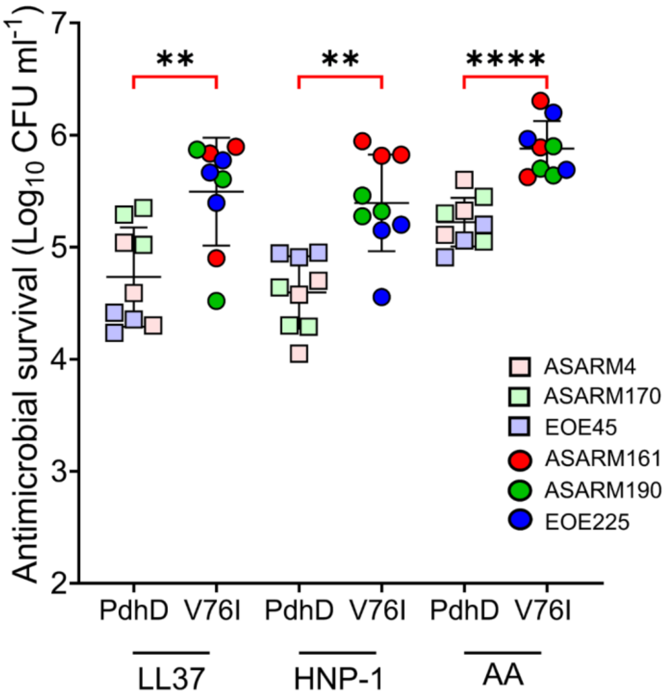

**Figure S3 – Growth kinetics of CC30 clinical strains in TSB (A) and RPMI-C (B).** Growth curves of CC30 clinical strains with wild type PdhD (ASARM4, ASARM170 and EOE45) or PdhD V76I (ASARM161, ASARM190 and EOE225). No differences in growth exhibited between strains expressing PdhD V76I or wild type PdhD. For growth kinetics points represent the average of three independent experiments with a minimum of three technical replicates per experiment. Readings ($OD_{600nm}$) measured with a Tecan PRO200 infinite plate reader.

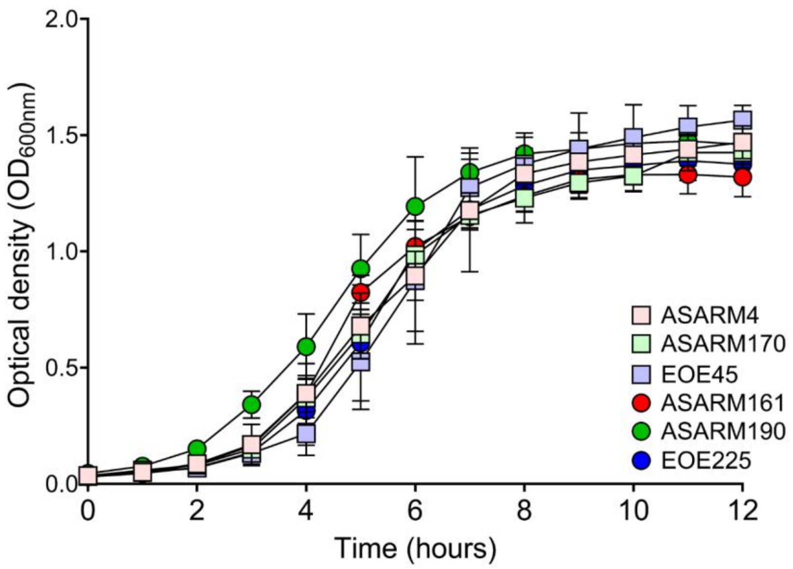

**Table S1 – Minimum inhibitory concentration assays for isogenic JE2 and SH1000 mutants with associated complements.** Complementation with PdhD V76I results in increased resistance to gentamicin and $H_2O_2$ when compared to wild type restoration (highlighted in yellow). Complementation with both *pdhD* and *pdhD* V76I restores daptomycin and cloxacillin resistance to wild type levels, with no enhancement seen for expression of PdhD V76I. MICs conducted by broth microdilution in flatbottomed 96-well plates using cation adjusted Mueller-Hinton broth (MHB). For daptomycin MICs MHB was modified with an additional 1.25 mM of calcium chloride. Values represent median of three repeats.

| | **Antimicrobial** | | | | |
|---|---|---|---|---|---|
| **Strain** | **Gentamicin** | **$H_2O_2$** | **Daptomycin** | **Cloxacillin** | **Teicoplanin** |
| JE2 | 0.5 | 4 | 0.5 | 1 | 1 |
| *pdhD*::tn | 4 | 2 | 0.125 | 0.125 | 1 |
| JE2 pEmpty | 0.5 | 4 | 0.5 | 1 | 1 |
| *pdhD*::tn pEMPTY | 4 | 1 | 0.125 | 0.125 | 1 |
| *pdhD*::tn p*pdhD* | 0.5 | 4 | 0.5 | 1 | 1 |
| *pdhD*::tn pSNP | 2 | 16 | 0.5 | 1 | 1 |
| SH1000 | 1 | 8 | 0.5 | 0.5 | 1 |
| *pdhD*::tn | 4 | 2 | 0.25 | 0.125 | 1 |
| SH1000 pEmpty | 0.5 | 8 | 0.5 | 0.5 | 1 |
| *pdhD*::tn pEMPTY | 4 | 2 | 0.25 | 0.125 | 1 |
| *pdhD*::tn p*pdhD* | 0.5 | 8 | 0.5 | 0.5 | 1 |
| *pdhD*::tn pSNP | 2 | 16 | 0.5 | 0.5 | 1 |

**Table S2 – Kinetic parameters assessing activity of PdhD vs PdhD V76I.** DLD assays assessed using variable concentration of $NAD^+$ and DHL whilst DP assay assessed using variable concentrations of NADH and MTT. Maximal reaction rate ($V_{max}$), substrate affinity ($K_m$), substrate molecule turnover per second ($K_{cat}$) and catalytic efficiency ($K_{cat}/K_m$) calculated and compared to wild type PdhD kinetic parameters. Kinetic parameters calculated using data presented in figure 4. Significance determined as *<0.05, **<0.01, ***<0.01 calculated by t-test

| Substrate | Parameter | Variant | |
|---|---|---|---|
| | | PdhD | Val76Iso |
| **NAD+** | $V_{max}$ (μM min$^{-1}$) | 348.2±21.3 | 331.1±23.5 |
| | $K_m$ (μM) | 3.184±0.82 | 4.045±1.1 |
| | $K_{cat}$ (s$^{-1}$) | 116.1±3.78 | 110.4±5.14 |
| | $K_{cat}/K_m$ (μM s$^{-1}$) | 36.46±9.55 | 27.29±6.64* |
| **Dihydrolipoic acid (DHL)** | $V_{max}$ (μM min$^{-1}$) | 182.8±8.95 | 191.6±7.35 |
| | $K_m$ (μM) | 1.548±0.327 | 2.487±0.597* |
| | $K_{cat}$ (s$^{-1}$) | 60.94±2.99 | 63.86±2.45 |
| | $K_{cat}/K_m$ (μM s$^{-1}$) | 39.36±6.73 | 25.68±5.1* |
| **NADH** | $V_{max}$ (μM min$^{-1}$) | 277.4±8.1 | 340.5±11.5*** |
| | $K_m$ (μM) | 0.3291±0.042 | 0.3032±0.046 |
| | $K_{cat}$ (s$^{-1}$) | 92.48±2.705 | 113.5±3.85** |
| | $K_{cat}/K_m$ (μM s$^{-1}$) | 280.93±27.15 | 374.34±43.7* |
| **MTT** | $V_{max}$ (μM min$^{-1}$) | 202.6±18.3 | 325.6±26.1** |
| | $K_m$ (μM) | 0.7169±0.23 | 0.8218±0.14 |
| | $K_{cat}$ (s$^{-1}$) | 67.54±6.11 | 108.5±8.7** |
| | $K_{cat}/K_m$ (μM s$^{-1}$) | 94.2±22.1 | 129.58±12.26* |

**Figure S4 – Total NAD/NADH pool in JE2 mutant and complement strains in TSB (A) and TSB-Serum (B).** Total NAD/NADH percentage normalised to JE2 wild type. Loss of *pdhD* results in a 45 and 66.5% reduction of the NAD/NADH pool when cultured in TSB and TSB-serum respectively. Complementation with wild type or PdhD V76I restores the NAD/NADH pool to wild type levels in both medias. No difference is exhibited between complementation plasmids, indicating that the altered NADH: $NAD^+$ ratios exhibited by PdhD V76I are not due to changes in total abundance of co-enzyme. Points represent individual experiments averaged from two technical replicates. Lines indicate the mean of these experiments with error bars denoting standard deviation. Statistical significance determined as *<0.05, **<0.01, ***<0.001, and ****<0.0001 by one-way ANOVA with Dunnett's T3 post-hoc test.

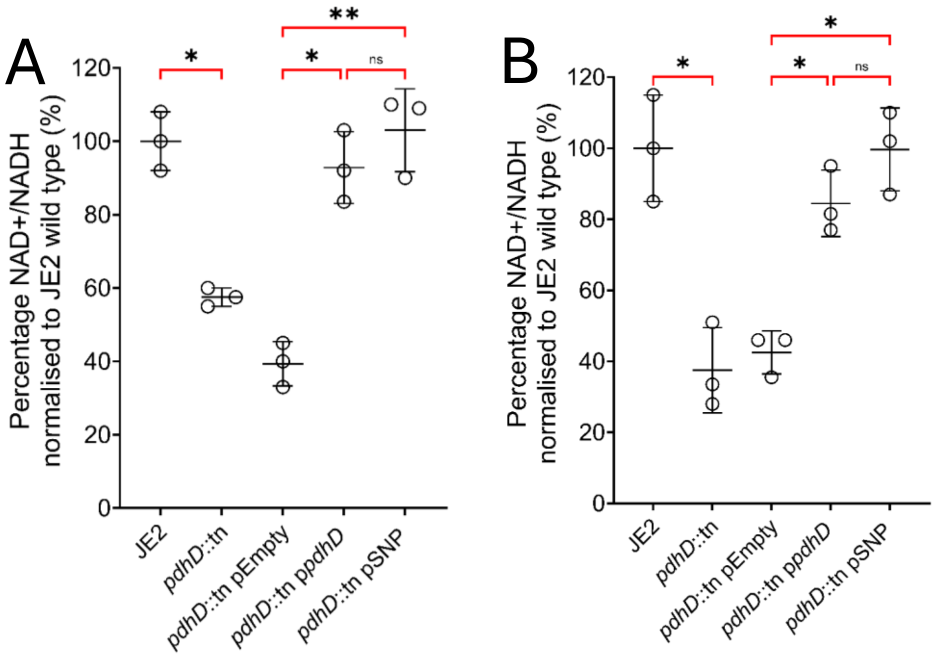

**Figure S5 - PdhD V76I mutation is associated with reduced haemolysis.** Loss of *pdhD* leads to reduced haemolysis of human erythrocytes with *agrA*::tn acting as a control. Complementation with wild type *pdhD* restores haemolysis to wild type levels, whilst restoration with PdhD V76I only partially restores toxicity. Points represent individual experiments with a minimum of 5 technical replicates; error bars represent standard deviation. Statistical significance determined as *<0.05, **<0.01, ***<0.001, by one-way ANOVA with Dunnett's T3 post-hoc test.

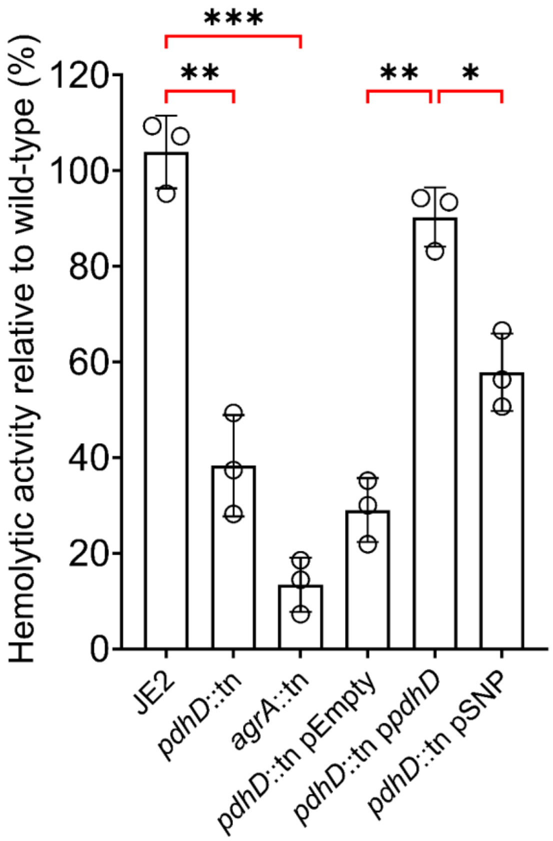

**Figure S6 – Loss of *ndhC* phenocopies PdhD V76I increased resistance to gentamicin and hydrogen peroxide.** Mutation of *ndhC* results in reduced sensitivity to both gentamicin and hydrogen peroxide due to reduced metabolic activity. Complementation restores sensitivity to both antimicrobials. Expression of the PdhD V76I mutation phenocopies loss of *ndhC*, with increased resistance to both gentamicin and hydrogen peroxide. This is consistent with expression of PdhD V76I reducing the ability of NdhC to process NADH, leading to reduced metabolic activity and enhanced resistance. Gentamicin and hydrogen peroxide killing assays conducted with an initial inoculum of $1 \times 10^7$ cells. Gentamicin killing conducted in TSB at 5 µg $ml^{-1}$ for 2 hours, hydrogen peroxide killing conducted in PBS at 20 mM for 90 minutes. Points represent individual experiments with a minimum of three technical replicates; bars represent the mean values with error bars standard deviation. Significance determined as *<0.05, **<0.01 calculated by One way ANOVA with Sidak's multiple comparisons test.

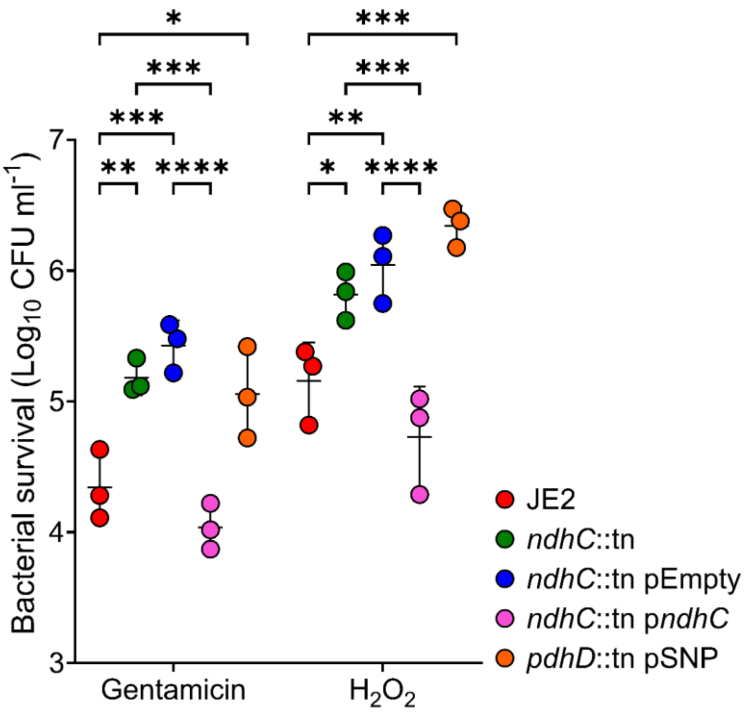
